# Peripheral blood DNA methylation partially reports on immune cell type composition in the frontal brain

**DOI:** 10.1101/2025.08.25.672204

**Authors:** Mandy Meijer, Maggie Po Yuan Fu, Erick Isaac Navarro-Delgado, Hannah-Ruth Engelbrecht, Gustavo Turecki, Meingold Hiu-ming Chan, Michael Steffen Kobor

**Affiliations:** Department of Medical Genetics, Faculty of Medicine, University of British Columbia, Vancouver, BC, Canada; British Columbia Children’s Hospital Research Institute, University of British Columbia, Vancouver, BC, Canada; Centre for Molecular Medicine and Therapeutics, University of British Columbia, Vancouver, BC, Canada; Edwin S. H. Leong Centre for Healthy Aging, Faculty of Medicine, University of British Columbia, Vancouver, BC, Canada; Department of Psychiatry, McGill Group for Suicide Studies, Douglas Mental Health University Institute, McGill University, Montreal, QC, Canada; Department of Human Development and Family Science, Purdue University, West Lafayette, IN, USA

**Keywords:** **<u>Keywords</u>**: DNA methylation, brain, epigenome-wide association study, machine learning, epigenetic age acceleration, NK cells, oligodendrocytes, astrocytes, microglia

## Abstract

Most existing DNA methylation (DNAm) studies have used peripheral surrogate tissues to research molecular mechanisms underlying brain disorders and diseases. Initial studies comparing brain to blood primarily at the individual CpG level analysis have generally pointed to a limited overlap of epigenetic patterns, consistent with DNAm being largely tissue- and even cell type-specific. Expanding on these studies, we employed a more complex analysis strategy aimed to 1) identify single DNAm sites associated with deconvolution-estimated brain cell type proportions, the principal measure in this study, in both the frontal brain and peripheral blood, 2) combine blood DNAm sites to predict brain cell type proportions through multivariate models, and 3) examine the association of blood DNAm, age, and epigenetic age acceleration (EAA) on brain cell type proportions. Epigenome-wide association studies for seven brain cell type proportions in matched frontal brain and peripheral blood samples (n=104) revealed that ∼10% of brain cell type-associated DNAm sites had correlating DNAm levels in peripheral blood (p<0.05). However, only three peripheral blood DNAm sites were significantly associated with endothelial and stromal brain cell type proportions (adjusted p<0.05). Brain cell type proportion predictions trained with machine learning approaches using peripheral blood DNAm showed the strongest, although still modest correlations with microglia proportions estimated through cell deconvolution using brain DNAm. Further, deconvolution-estimated blood immune cell type proportions showed a nominally significant association with estimated brain stromal cell proportions, driven primarily by NK cells; however, this association was dependent on chronological age and did not survive age-residualization. Lastly, brain EAA was not associated with brain and blood cell type proportions (adjusted p<0.05). Collectively, these results suggested that in the context of broad tissue-specificity of DNAm patterns, DNAm levels in peripheral blood might actually inform on some immune brain cell type proportions. The correlations between DNAm profiles specific to immune cell types in blood and brain were consistent with a potential link between peripheral immune and central nervous system immune functions.

## 1. Background

Brain-related disorders and diseases are common and have high lifetime prevalences. For instance, one in eight people in the world live with a psychiatric disorder (1), and the lifetime risk of receiving a diagnosis of Alzheimer’s disease (AD) or Parkinson’s disease (PD) is up to one in five (2–4). Furthermore, psychiatric and neurodegenerative conditions are often co-occurring (5–8). These brain-related conditions, including neurodevelopmental disorders, psychiatric disorders, and neurodegenerative diseases, are multifactorial, involving both genetic as well as environmental factors in their development and persistence (9–13). Considerable knowledge about the molecular underpinnings of many brain-related diseases is lacking, and effective treatments for most of these conditions are not available yet.

Previous studies on postmortem brain samples have identified potential roles of specific brain cell types in different brain-related conditions, likely related to their specific functions. Neurons are the main brain cells that receive and send information; glial cells have a supporting role in the central nervous system: microglia and astrocytes are important effector cells in the central immune system (14), whereas the main role of oligodendrocytes is to myelinate neurons (15). The combination of differences in both the proportions of brain cell types and their actual cellular function in brain-related diseases is found for a number of cell types. For example, alterations in microglia proportions and microglial gene expression levels have been observed for autism spectrum disorder, schizophrenia, depression, and AD (16–20). Similar differences in gene expression levels have been observed for astrocytes in depression (17), and for gene expression levels in neuronal cells in autism spectrum disorder, schizophrenia, and PD, as well as cellular functionality and morphological differences in depression (16, 21–23). From a related, yet conceptually different perspective based on genome-, transcriptome-, and epigenome-wide association studies, genetic and epigenetic loci associated with brain-related conditions are enriched in different brain cell types: schizophrenia has been shown to be associated with median spiny neurons (24) and GABA- and glutamatergic neurons (25). Similar enrichment studies for ADHD have shown associations with excitatory neurons and astrocytes (26, 27), while PD has been associated with oligodendrocytes through genetic enrichment in cell-specific expression datasets (25). Collectively, this emerging evidence provides compelling examples that different brain-related conditions are linked to changes in a distinct palette of brain cell type proportions and functions. This then raises the opportunity that studying brain cell type composition more broadly might contribute an important layer to a better understanding of brain disease etiology.

Cell type identity, function and regulation are modulated, in part, by potentially heritable modifications of the chromatin landscape, often referred to as epigenetics. Of these, at least in human populations, DNA methylation (DNAm) is best understood. DNAm patterns are tissue- and even cell type-specific (28), and tend to be more stable than transcriptomics signatures (29–31). However, most DNAm-association studies on brain-related outcomes are performed in surrogate tissues such as blood. In living humans, the single most important reason for the choice of peripheral samples over brain samples is practicality related to less invasive collection protocols (32). That said, given the major relation of DNAm patterns with tissue and cellular identify, it is clear that peripheral blood samples might not fully report on brain DNAm. This has been substantiated by, previous studies documenting that the relationship between DNAm in peripheral surrogate tissue and brain issue is rather complex. On the one hand, DNAm levels show statistically significant global correlations between peripheral blood and brain tissue across all measured DNAm sites (Pearson’s r=0.860 and r=0.892), although this might be skewed by the inclusion of non-variable DNAm sites (33–36). On the other hand, at the level of individual peripheral DNAm sites, only 6-11% showed moderate to high blood-brain DNAm level correlations (33–36). Despite the cell type and tissue-specificity of DNAm, the correlation of DNAm sites across tissues suggests that some peripheral blood DNAm sites can still be informative for brain DNAm levels, as documented to various degrees in previous studies. On that basis, single DNAm sites with high blood-brain correlation might lead towards informative biomarkers for brain-related phenotypes. Supporting this notion, epigenome-wide association studies have identified DNAm sites that are significantly associated with different brain disorders using peripheral samples (37–43).

It is possible that associations of blood-based DNAm loci with brain disorders might at least in part reflect the biological cross-talk and physical interaction of the peripheral and central nervous system (44). It has been shown recently that the body-brain axis signals to the brain about an emerging inflammatory response during neurodegenerative diseases and the ageing process. In turn, the brain can modulate the peripheral immune response, as well as immune infiltration in the central nervous system, (110, 111). The potential close nexus between brain and blood through the immune system is further supported by a high comorbidity and genetic correlation between brain disorders and autoimmune diseases (45–50), and many conditions are thought to have an immune system component, such as schizophrenia (51), bipolar disorder (52), ADHD (53), and PD (54). Thus, investigating the correlation between peripheral immune cell proportions and brain cell type proportions is an important avenue towards a better understanding of these disorders.

Alterations in immune functioning are not only associated with brain diseases, but also a notable part of the aging process. As such, age is often identified as an important risk factor related to the development of psychiatric disorders (e.g., depression (55)), and neurological disorders (e.g., dementias (2)). Studying the molecular markers of ageing can thus provide a useful avenue to interrogate brain-related conditions, which increase with ageing (56), and their associations with immune system functioning, which decreases with ageing (57). It is important to delineate the effects of ageing on brain cell function and proportions, which can be studied through the lens of biological ageing. Epigenetic age acceleration (EAA), a form of biological ageing, is the relative rate of ageing for an individual given their chronological age, estimated based on DNAm data (58–60). EAA has been associated with age-related brain diseases and traits (61–63), as well as multiple psychiatric disorders and neurodegenerative diseases (e.g., (61–71)). There are many factors potentially driving EAA. For example, it has been previously shown that differences in peripheral blood epigenetic age is associated with Natural Killer (NK) cell and T cell immunosenescence (72), and that oligodendrocytes are associated with increased epigenetic ageing in brain (73).

In summary, the exact underlying molecular mechanisms of many brain diseases are unknown, but changes in brain cell type proportions have been identified to potentially underlie different disorders and diseases. However, access to brain samples is limited, hampering research into the molecular mechanisms of brain-related traits in humans. Because most studies are relying on peripheral blood samples, the biological interpretation of the existing correlation between peripheral blood and brain DNAm could be better understood when taking cell type proportions into account. In addition to relying on single DNAm sites, multivariate analyses could also reveal new informative patterns linking blood DNAm and brain cell types that could be left undiscovered by single site approaches. Lastly, it is important to establish the relation to age in these correlations. Therefore, the overarching goal of this study was to investigate whether DNAm in peripheral blood can inform brain cell type-specific epigenetic signals. Focusing the analysis of the complex information content of DNAm derived from surrogate tissues and brain, this mapping exercise adds a different dimension to the existing literature on studying brain-related diseases and brain health in general. Specifically, we investigated brain DNAm in the frontal cortex, as this brain region is affected in many brain-related conditions, including ADHD, schizophrenia, and bipolar disorder (74). We aimed to 1) identify single DNAm sites associated with deconvolution-estimated brain cell type proportions in both the frontal brain and peripheral blood; 2) combine blood-based DNAm sites to inform brain cell type proportions through multivariate models, and 3) identify the association of chronological age and biological brain ageing on brain cell type proportions.

## 2. Methods

### 2.1 Samples

Publicly available datasets containing paired blood and brain DNA methylation (DNAm) were used (GSE214901; GSE95049; GSE111165; GSE59685) (**Table S1**). Detailed descriptions of the individual datasets have been published previously (33–36). Two datasets included live brain tissue samples from patients undergoing neurosurgery for clinical purposes (GSE214901; GSE95049), and two datasets included postmortem brain tissue dissected at autopsy (GSE111165; GSE59685). All brain samples taken from the frontal brain were selected. All brain samples that had matched blood samples from the same individual were retained. Of these, N=30 (GSE214901; GSE95049; GSE111165) blood and brain samples were collected at the same time point, and N=74 (GSE59685) blood and brain samples were collected with a mean delta time of 3.83 years (**Table S1**). When one dataset included more than one frontal brain sample from the same individual, one random sample was selected. In total, we included 104 matched frontal brain and peripheral blood samples from the same individuals in the current study.

### 2.2 DNA methylation

Our work relied on existing DNAm data; no new DNAm arrays were generated. In general, all the utilized studies used DNA extracted from the whole blood and brain samples as previously described (33–36). DNAm was assessed by either the Infinium® HumanMethylation450 BeadChip (450K array; Illumina, San Diego, CA, USA) or the Infinium® MethylationEPIC BeadChip (EPIC array; Illumina), and only probes overlapping between the two arrays were retained. Raw data, both for each dataset and tissue type, were processed separately in R v4.2.4. For dataset 1, 2, and 4 (**Table S1**), raw iDAT files were available and processed, for dataset 3, raw beta values were available and processed. A complete overview of DNAm preprocessing and quality control (QC) can be found in the **Supplementary methods**. In total, 366,570 and 370,721 DNAm sites were retained in blood and brain for subsequent analysis, respectively, after extensive QC.

### 2.3 Cell type deconvolution

Brain cell type proportions were estimated through epigenomic deconvolution in the *HiBED* (Hierarchical Brain Extended Deconvolution) v0.99.6 R package (75). Proportions were estimated for astrocytes, endothelial cells, GABA- and glutamatergic neurons, microglia, oligodendrocytes, and stromal cells. It should be noted that the endothelial and stromal references have been derived from newborn cord tissue and blood in the HiBED panel. The HiBED reference panel consists of two layers with categories (L1, L2A, L2B, and L2C), containing 621 probes in total. In our dataset, 580 probes were present: 75 out of 81 for L1, 174 out of 183 for L2A, 219 out of 237 for L2B, and 112 out of 120 for L2C.

Blood cell type proportions were estimated for twelve different immune cell types based on the IDentifying Optimal Libraries (IDOL) extended reference dataset (76): basophils, eosinophils, neutrophils, monocytes, B naïve cells, B memory cells, CD4^+^ naïve cells, CD4^+^ memory cells, CD8^+^ naïve cells, CD8^+^ memory cells, NK cells, and T regulatory cells. Betas from dataset 1 through 4 were quantile normalized with the betas present in the reference panels, and probes used to estimate cell type proportions were selected based on t-tests comparing each cell type against others (e.g., neutrophils vs non-neutrophils), with a p-value cut-off of 0.05, and minimum delta beta of 0.05. One hundred probes were selected for each cell type estimated, which is the default for most DNAm-based deconvolution algorithms (77, 109). Cell type proportions were estimated using constrained projection (76, 77). Interquartile ranges (IQR) for the probes selected for blood cell type proportion estimations were calculated.

### 2.4 Epigenome-wide association meta-analyses

To identify whether single DNAm sites in blood inform brain cell type proportions, we performed epigenome-wide association studies (EWAS) for each brain cell type, separately by using linear regression models corrected for age, sex and cell type proportions. We conducted such EWAS in both blood as well as brain samples, separately. To make results comparable across datasets 1 through 4 and across tissues (i.e., brain and blood), we also corrected for biological variability likely attributable to tissue-specific cell type proportions for the tissue involved in the EWAS through surrogate variable analysis (SVA). For each statistical model we selected the optimal number of surrogate variables needed to account for variation in the data based on permutation. An overview of the number of surrogate variables for each analysis can be found in **Table S2**. Results for each brain cell type in each tissue (i.e., blood and brain) were sample size-based meta-analyzed using METAL (78). FDR-adjusted p-values were calculated over all seven cell types tested per tissue (e.g., number of tests for brain = 7*370,721; number of tests for blood = 7*366,570), and DNAm sites with an Benjamini-Hochberg (BH) adjusted p-value<0.05 were considered statistically significant in the meta-EWAS. Furthermore, DNAm sites were considered statistically significant if the heterogeneity test p-value was > 0.05, (meaning there was no significant heterogeneity between datasets) *and* if direction of effect was consistent across all four datasets (**Figure S1**). Inflation factors for each EWAS were calculated as the square root of the expected median of a chi-squared statistics distribution divided by the median of the chi-statistics with one degree of freedom. To determine whether percentages of correlations between blood and brain for the significant DNAm sites were more or less than expected by chance, we performed a permutation analysis. Specifically, brain DNAm sites were randomly shuffled ten times and analyses were repeated. While ten permutations is modest, the resulting null distribution was highly stable (SD=0.15%), supporting its use for inference here. Where the findings from the observed analysis did not overlap with this null distribution, results were considered likely to reflect biological signal rather than statistical artifact.

### 2.5 Multivariate brain cell type proportion predictors based on blood DNAm data

To assess whether a multivariate predictor of brain cell type proportions could be created out of DNAm sites in peripheral blood, Ridge regression was performed on a training dataset and validated in a testing dataset. The training superset consisted of datasets 1, 2, and 3. Prior to Ridge regression, the training superset was renormalized together with *dasen* and *ComBat*. Dataset 4 was reserved as test dataset and renormalized by itself. Additionally, to reduce dimensionality, Variable Methylated Regions (VMRs) were generated using the *RAMEN* R package (79) using the default parameters. A Random Forest (RF) model was then trained using the *ranger* R v0.14.1 package with 500 trees (80). See **Supplementary methods** for more details on the Ridge regression, VMRs and RF.

### 2.6 Principal component analysis

To assess whether DNAm sites in peripheral blood with high blood-brain correlation can inform brain cell types, we used the 429 probes from the brain cell type reference panel that were present in both the frontal brain and blood supersets. We used this specific set of probes as these have been identified to distinguish between different brain cell types in datasets independent of ours (75). We calculated statistically significant correlations (p<0.05) between these DNAm sites in both blood and brain. We used these, rather than the DNAm sites significantly associated with brain cell type proportions in brain tissue, as the reference probes reflect a hierarchical granularity (i.e., containing probes informative for both glial cells (Layer 1) and e.g., microglia (Layer 2), and are derived from single-cell measures (75). The DNAm sites were clustered by cell type (e.g., for microglia all sites annotated to either microglia or glial cells were clustered), and a principal component analysis (PCA) was performed on the derived DNAm sites in both brain and blood.

### 2.7 Epigenetic ageing in the brain

Epigenetic frontal brain age was calculated with the Cortical Clock (60) and epigenetic age acceleration (EAA) was calculated by residualizing predicted epigenetic brain age on chronological age. Samples with an EAA > |25| were removed, which eliminated one potential outlier (EAA=50). The median chronological age of the individuals used in this analysis was 76.76 years.

## 3. Results

### 3.1 Dataset description

We combined four publicly available datasets of brain and blood DNAm (GSE214901; GSE95049; GSE111165; GSE59685), and retained in total 104 frontal cortex and blood samples that were matched from the same individuals (**Table S1**). The mean age of brain and blood samples at collection were 76.76 years and 74.04 years, respectively (Standard Deviation (SD) = 21.62 and 20.30 years, respectively), where the median age difference for collection was 2.00 years (range: 0-16 years). No effect of age differences was observed in overall DNAm patterns between the four cohorts (**Figure S2**). In total, 44% of the individuals were male. Using this combined dataset, we tested whether DNAm in peripheral blood can inform brain cell type-specific epigenetic patterns, using three distinct yet overlapping broad approaches, ranging from association studies to machine learning-based predictors (**Figure 1**).

**Figure 1.**
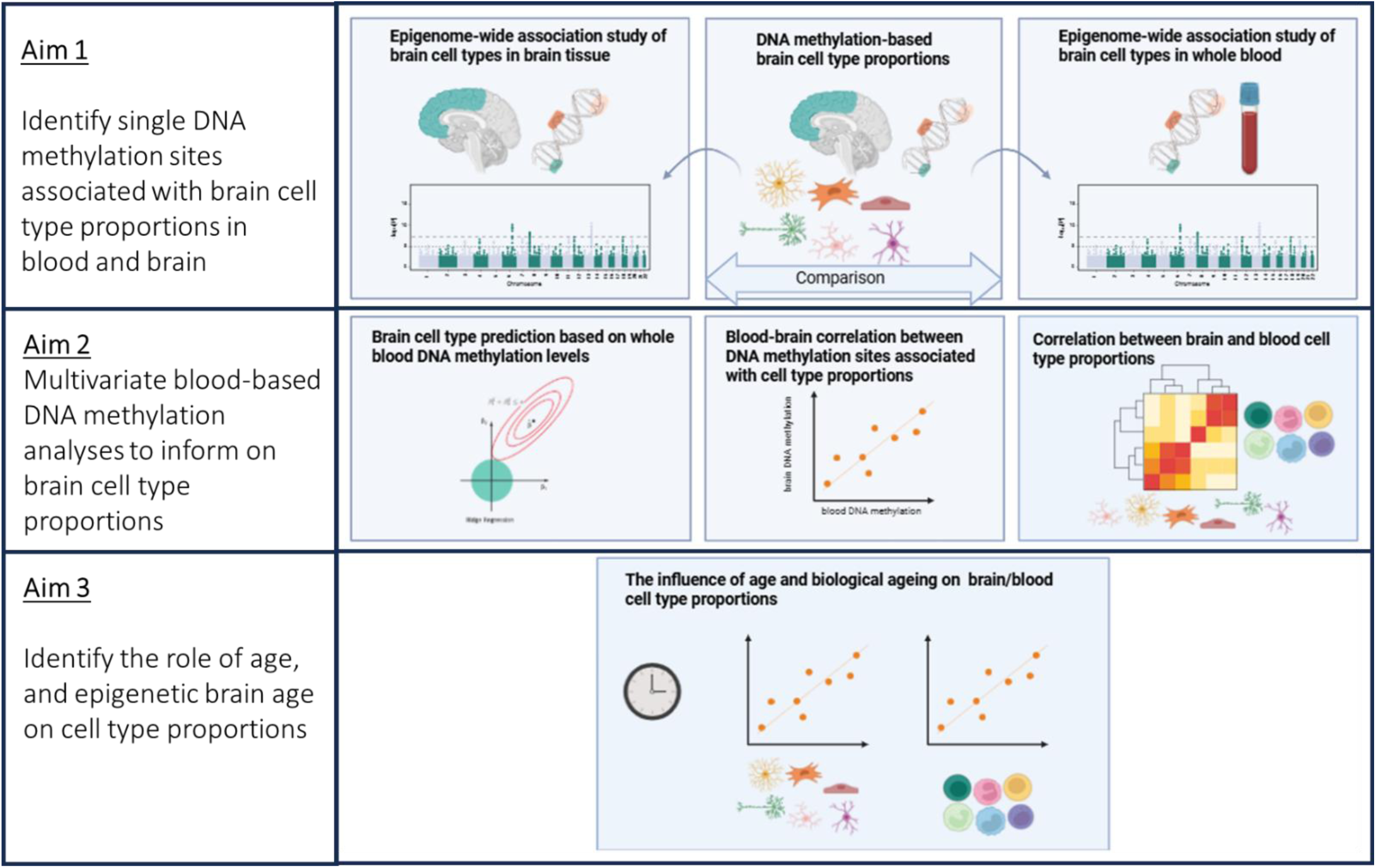
Study overview. Aim 1: DNA methylation (DNAm) was assessed in paired frontal cortex and blood samples (upper row, left and right panels). Brain cell type proportions (astrocytes, endothelial cells, stromal cells, oligodendrocytes, GABAergic neurons, glutamatergic neurons and microglia) were estimated based on epigenomic deconvolution in frontal cortex samples (upper row, middle panel). Epigenome-wide association studies (EWAS) for each brain cell type proportion were performed in both brain and blood samples (upper row, left and right panels). The results of these EWAS were compared across tissues. Aim 2: Then, the dataset was split in a training and testing dataset and all DNAm sites in blood were used to predict brain cell type compositions through Ridge regression (middle row, left panel). We also assessed whether a subset of DNAm sites with highly correlated blood-brain DNAm levels were associated with brain cell type proportions (middle row, middle panel). Because of the consistent immune brain cell types-blood DNAm associations, blood immune cell type proportions were estimated, and tested for correlations with estimated brain cell types (middle row, right panel). Aim 3: Then, the influence of age on these correlations was assessed (bottom row). Epigenetic age acceleration in the cortex was estimated based on cortical epigenetic clocks, and the association of different brain and blood cell types on biological brain ageing was assessed.

As the foundation for our study, we first used established procedures for bioinformatic deconvolution of DNAm data derived from the frontal brain to estimate proportions of seven different cell types in all 104 samples. As expected, oligodendrocytes were the most abundant cell type present in the brain samples (mean = 38.15%, inter-quartile range (IQR) = 9.36%), followed by glutamatergic neurons (mean = 25.72%, IQR = 6.35) (**Figure 2**). Furthermore, their levels were highly negatively correlated with each other (Pearson’s r = -0.91, p < 2.2*10^-16^). This correlation was not entirely unexpected since cell type proportions are compositional (e.g., they sum up to 1), and oligodendrocytes and glutamatergic neurons have the highest proportion among cell types. We noted a considerable variability for both of these deconvolution-estimated cell types in each of the individual datasets, whereas the other five cell types had a tighter distribution, perhaps owing to their overall lower proportions. All datasets showed similar cell type proportion levels, irrespective of consisting of live or postmortem tissue. These deconvolution-estimated cell type proportions were the principal input of the subsequent analyses.

**Figure 2.**
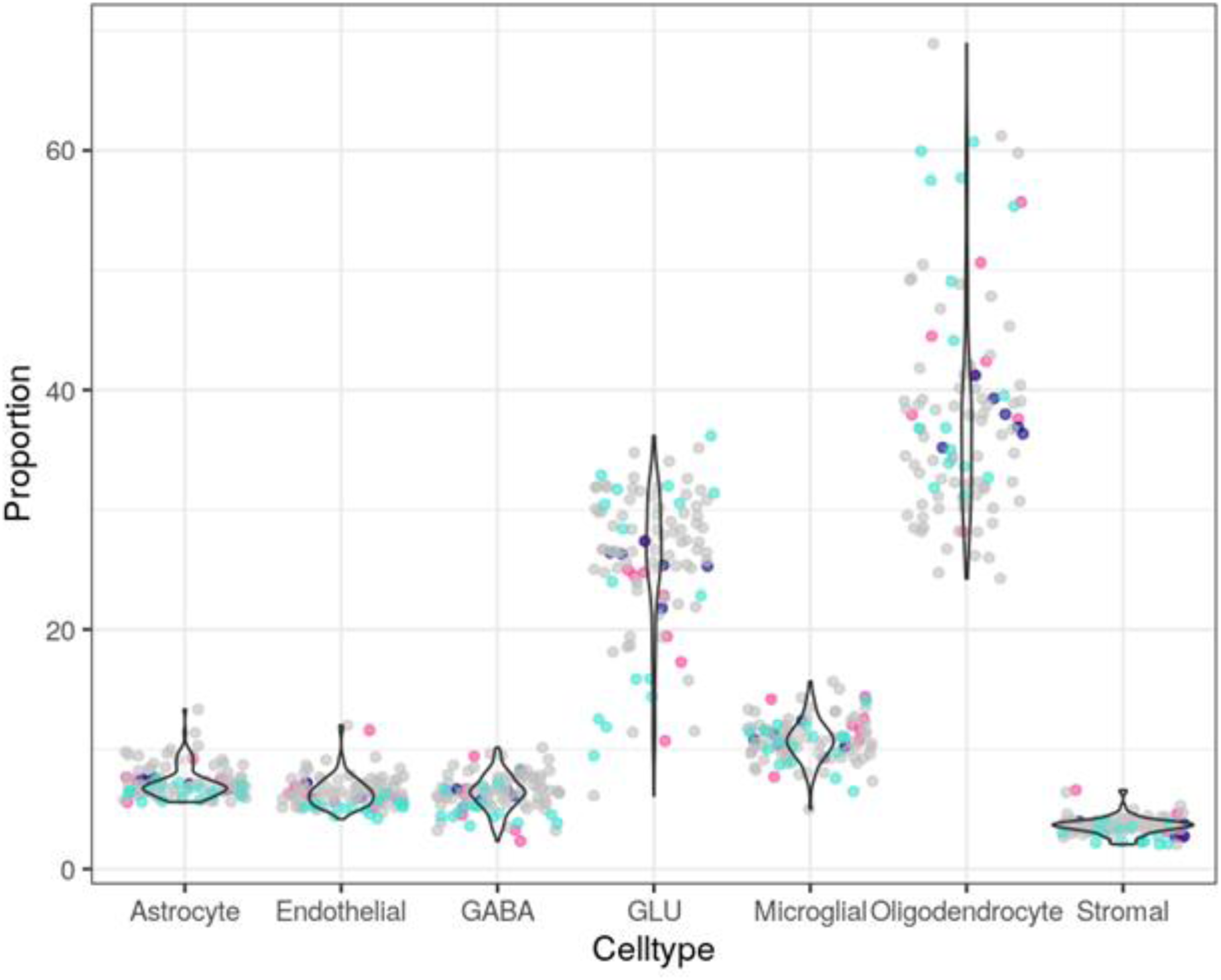
Oligodendrocyte proportions were most abundant and variable in the frontal brain. Brain cell type proportions (astrocytes, endothelial cells, GABAergic neurons, glutamatergic neurons, microglial, oligodendrocytes, and stromal cells) in the frontal brain were estimated in the combined datasets based on the epigenomic deconvolution tool HiBED. Different colours indicate the individual sub-datasets: Dataset 1 = violet red; dataset 2 = navy blue; dataset 3 = grey; dataset 4 = turquoise blue.

### 3.2 DNA methylation associations with brain cell type proportions were tissue-specific

Given the role of DNAm in the establishment and maintenance of cellular identity, we next tested which individual DNAm sites in bulk brain DNAm profiles were associated with brain cell type proportions in the frontal cortex specifically. To account for heterogeneity within datasets and to identify DNAm sites with consistent direction of effect across datasets, we stratified by dataset and performed a meta-EWAS (**Figure 3**, **Table 1, Figure S1, 4-7, Table S3**). In total, 370,721 DNAm sites were retained for brain tissue after extensive quality control, which included the CpGs that were used to estimate the brain cell type proportions. Inflation of test-statistics could be explained by the dependency between brain DNAm and brain cell type proportions, leading to many significant associations (**Figure S4**). In the frontal brain, after removal of DNAm sites with inconsistent direction of effect for the association with brain cell types across the four different datasets, the number of brain DNAm sites significantly associated with brain cell type proportions ranged from seven for stromal cells to 54,425 for oligodendrocytes, however, at the level of individual CpG sites, effect sizes of these associations were similar (**Figure S7**). We also identified 94 GABA-DNAm associations, 1,623 endothelial cell-DNAm associations, 7,450 astrocyte-DNAm associations, 8,391 microglia- DNAm associations, and 36,217 GLU-DNAm associations. There was a considerable amount of overlap between GLU-DNAm and oligodendrocyte-DNAm associations, likely driven by the strong correlation between these cell type proportions due to the compositional nature of the cell type proportions (**Figure 3**). Of all the significant DNAm-brain cell type proportion associations (adjusted p-value<0.05), 77% of associations with GABAergic neurons had a positive regression coefficient in the meta-EWAS, whereas the associations for astrocytes, glutamatergic neurons, and stromal cells primarily had negative regression coefficients (58%, 64%, and 71%, respectively).

**Figure 3.**
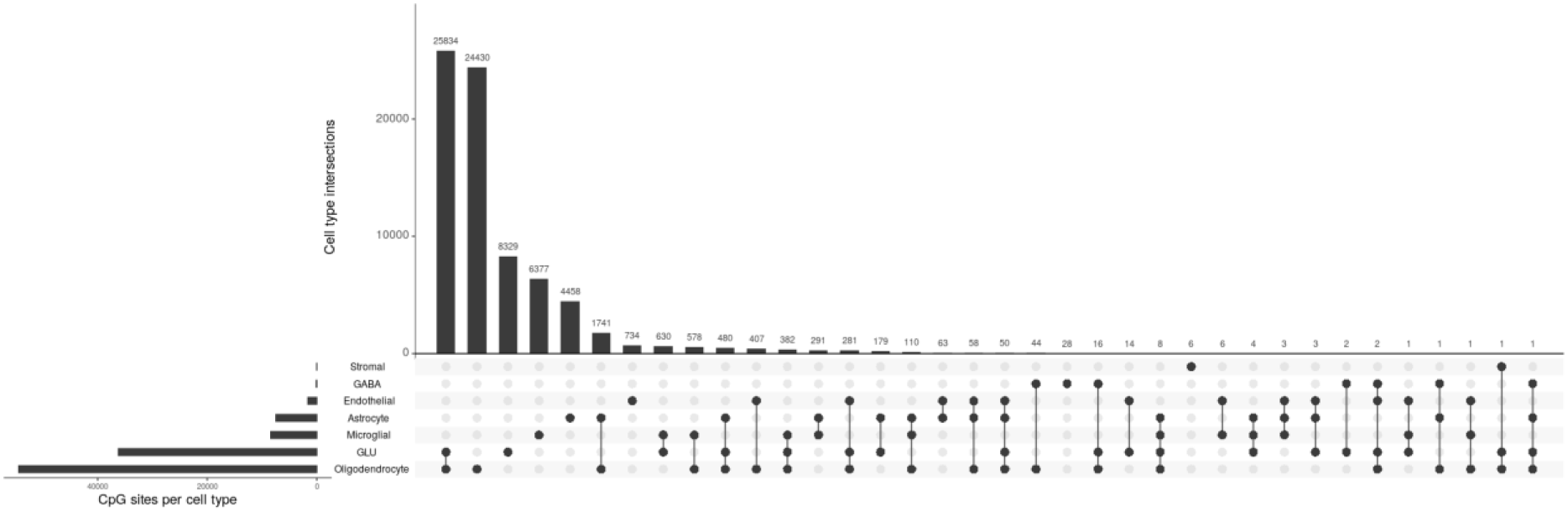
A subset of brain DNA methylation associations with distinct brain cell types was overlapping with each other. A meta-EWAS from bulk frontal cortex-derived DNA methylation profiles was performed for seven different brain cell type proportions. Each cell type is represented as a row. On the left, the horizontal bar plot indicates the number of significantly associated DNA methylation sites per cell type as identified by the meta-EWAS. On the top, the vertical bar plot shows the intersections (i.e., significantly associated DNA methylation sites which are overlapping) per combination of cell types. The filled black dots connected by lines in the rows indicate which combinations of cell types are displayed, with single dots reflecting the unique DNA methylation sites significantly associated with that specific cell type.

**Table 1.** Overview of significant findings for each EWAS for brain cell type proportion in frontal brain tissue.

|  |  | Brain DNA methylation |  |  |  |  | Brain & blood DNA methylation | Blood DNA methylation |
| --- | --- | --- | --- | --- | --- | --- | --- | --- |
|  | Variability of cell type in superset (sample variance) | # of significant DNAm sites (FDR-adjusted) | # of significant DNAm sites after removal of heterogenous test-statistics | # of significant DNAm sites with consistent direction of effect | % of DNAm sites with positive association | # (%) of 580 DNAm sites of the epigenomic deconvolution reference panel in significant DNAm sites | % of DNAm sites with blood-brain correlation (% after FDR-correction) | # of FDR- adjusted significant DNAm sites |
| **Astrocytes** | 1.75 | 13,310 | 11,558 | 7,450 | 42% | 38 (6.55%) | 8.9% (2.13%) | 0 |
| **Endothelial cells** | 1.61 | 6,277 | 4,323 | 1,623 | 60% | 4 (0.69%) | 9.43% (2.09%) | 1 |
| **GABAergic neurons** | 2.27 | 110 | 104 | 94 | 77% | 5 (0.86%) | 8.51% (0%) | 0 |
| **Glutamatergic neurons** | 37.28 | 54,029 | 43,331 | 36,217 | 44% | 152 (26.21%) | 10.11% (2.70%) | 0 |
| **Microglia** | 3.22 | 11,918 | 11,072 | 8,391 | 38% | 54 (9.31%) | 12.53% (4.76%) | 0 |
| **Oligodendrocytes** | 80.15 | 80,903 | 68,396 | 54,425 | 48% | 206 (35.52%) | 10.35% (2.94%) | 0 |
| **Stromal** | 0.55 | 12 | 10 | 7 | 29% | 0 (0%) | 14.29% (0%) | 2 |

*a Variability correlated with the # of significant DNA methylation sites identified per cell type*

We next asked, as a sensitivity analysis, whether any of the DNAm sites that were included in the publicly available cell type deconvolution algorithm actually were associated with brain cell type proportions. Of the 580 DNAm sites HiBED brain deconvolution reference panel, 55% (n = 286) included in the reference panel for all brain cell types combined (**Table 1**), which is more than expected by chance (permutation p < 1*10^-5^). These 286 DNAm sites were in the top 10% most significant EWAS results for each individual cell type. These findings indicated that the DNAm sites needed for brain cell type deconvolution were indeed associated with brain cell type, yet that there were also many DNAm sites (99.62% of significant findings) not involved in the deconvolution algorithm that were associated with brain cell type proportions.

As the next step, to understand the potential information content provided by blood for brain cell type proportion, we tested the extent to which brain cell proportion-associated DNAm sites identified in brain were concordant in blood. Of all the significant DNAm associations with any brain cell type proportion, 10.42% had significantly correlating DNAm levels between matched frontal brain and blood samples (p<0.05; Pearson’s r: |0.21 – 0.99|). After FDR-correction, only 2.79% of these DNAm sites showed a significant correlation (adjusted p-value<0.05). This is slightly more than expected by chance as permutation analysis reveals that 7.39% (SD=0.15%) and 0% (SD=0.0059%) of random probes showed a correlation between blood and brain with both statistical cut-offs, respectively.

Moving beyond the initial set of brain DNAm sites associated with cell type proportions, we asked whether DNAm levels in peripheral blood were also associated with brain cell type proportions, and thus whether these sites could help inform brain cell type proportions in association with a brain phenotype based on EWAS in peripheral blood. For this, we again used a similar meta-EWAS approach as for the brain DNAm-specific associations. In total, 366,570 DNAm sites were retained for blood after extensive quality control. In contrast to the brain-specific analysis, we identified only a limited number of statistically significant blood DNAm associations with distinct brain cell type proportions (**Figure S4-5**, **Table S4**): two DNAm sites in blood were significantly associated with endothelial cells (|effect sizes| in individual cohorts all below 0.034, adjusted p-value < 0.05) and one DNAm sites was associated with stromal cells (|effect sizes| in individual cohorts all below 0.04, adjusted p-value < 0.05). The genes these DNAm sites are located in show high expression levels in the brain for both neuronal and glial cells, according to the Human Protein Atlas (proteinatlas.org, accessed on 17/07/2024). No DNAm sites from peripheral blood were associated with the other five predicted brain cell type proportions. To assess whether the significant associations were unique to blood or shared with the brain results, we compared the results obtained from the EWASs performed in blood and brain tissue: two out of the three blood-based DNAm sites showed blood-brain correlation (Pearson’s r: 0.398 - 0.900) (**Table S4**), but none of these DNAm sites were significantly associated with any brain cell type proportion in the EWAS performed on DNAm retrieved from the frontal cortex (**Table S3**). Indeed, and perhaps not surprisingly given the distinct nature of the two tissues, concordance in general between the effect sizes at the level of single DNAm sites retrieved from the meta-EWAS in blood and frontal brain tissue was low (**Figure S8**).

### 3.3 Predicted brain cell type proportions in peripheral blood showed the strongest correlation with deconvolution-estimated immune brain cell types

As an alternative to the limited information content offered by individual blood DNAm sites on brain cell proportions, we tested whether a more complex machine learning-based combination of DNAm sites might be more promising for this purpose. To develop a predictive model of brain cell type proportions in the frontal brain based on a combination of blood DNAm sites, we used Ridge regression on a training superset consisting of combined and renormalized datasets 1, 2 and 3, whereas dataset 4 was independently normalized and used as a testing dataset (**Table S1**). We chose this splitting of our datasets to ensure a set of samples with a diverse age range, especially for the training set. Using this approach, the root mean squared error (RMSE) of the model trained using Ridge regression for the different cell type percentages were the following: astrocytes = 1.503, endothelial = 0.643, GABA = 1.676, GLU = 9.597, microglia = 3.701, oligodendrocyte = 16.071, stromal cells = 0.157. Perhaps owing to our limited sample size, our model did not perform uniformly across all cell types. Specifically, we noted that errors tended to be large for the most abundant cell types (oligodendrocytes and glutamatergic neurons), which also showed the greatest absolute variability in proportions. RMSE values were smaller in absolute terms for less abundant cell types, although their often narrow proportion ranges make the interpretation more challenging. Nevertheless, some patterns emerged that may possibly point to physiological relevance. For example, Ridge-predicted brain cell type proportions from blood DNAm correlated nominally with deconvolution-estimated microglia, the brain’s resident immune cells (Pearson’s r = 0.525, unadjusted p-value = 0.0366, adjusted p-value = 0.133) (**Figure 4**). In contrast, even though we identified significant blood DNAm-endothelial and DNAm-stromal cell associations in the EWAS meta-analysis, and the RMSE of the Ridge regression was low [RMSE=0.157], all proportions were estimated to be 0 for endothelial and stromal cell types in the testing dataset.

**Figure 4.**
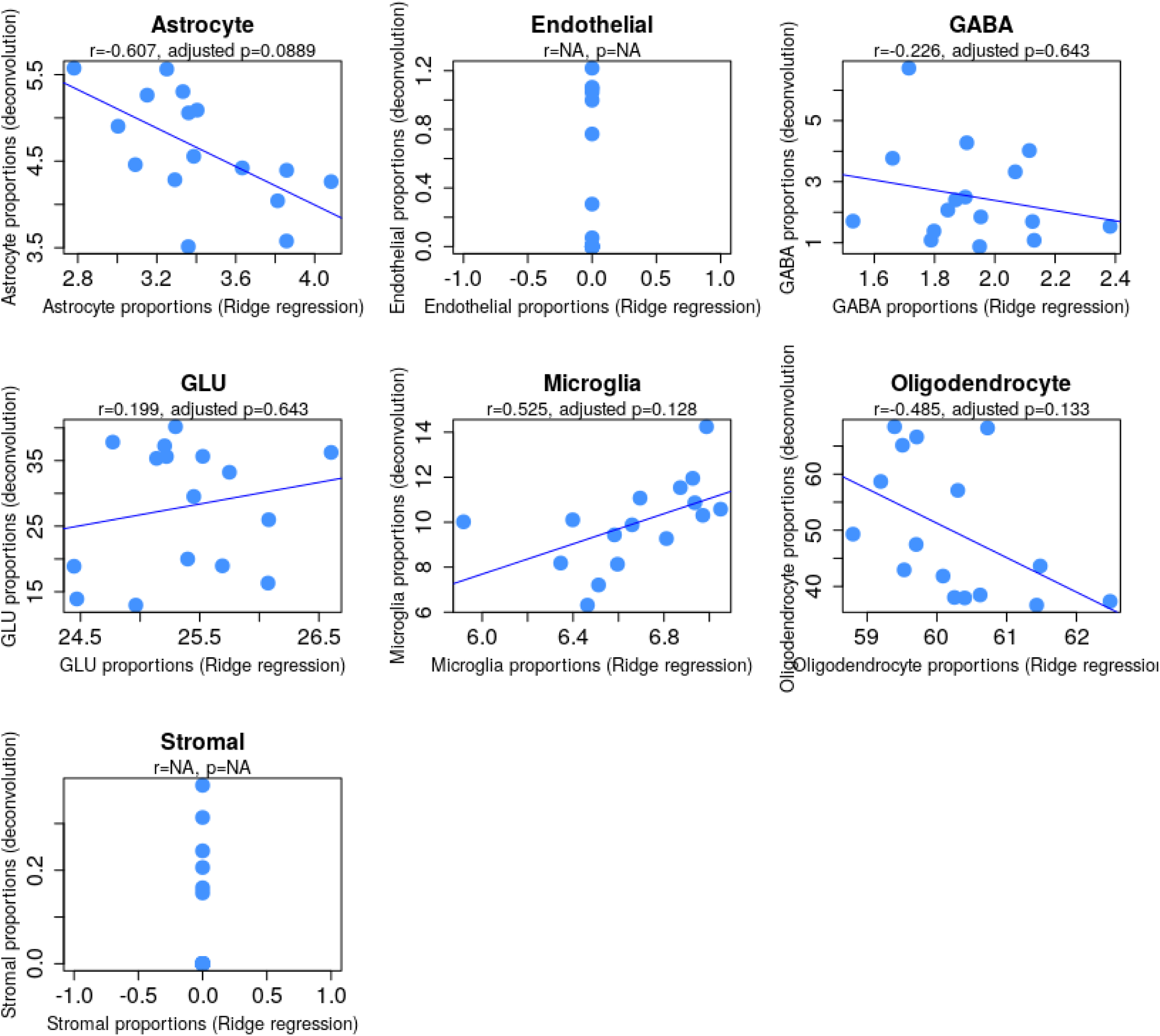
Brain immune cell type proportions showed the strongest associations with peripheral blood DNA methylation. Estimated brain cell type proportions based on epigenomic deconvolution in DNA methylation of frontal cortex samples is plotted on the y-axis. Brain cell type proportions estimated based on the optimal models from Ridge regression in blood are plotted the x-axis. Every dot represents a single sample from the renormalized test datasets 4 (**Table S1**). The Pearson’s correlation (r) and p-value (p) are given for every brain cell type.

While the performance of our initial model was moderate in predicting immune cell types, we considered potential improvements by reducing redundancy and increasing statistical power through the creation of variable methylated regions (VMRs), taking both the pairwise DNAm probe correlation and variability of DNAm probes into account. In addition, we included a Random Forest model for the VMRs to compare model performance. Given that the use of VMRs did not improve model performance for either Ridge regression (**Figure S9**) or Random Forest models (**Figure S10**), we performed a sensitivity analysis to determine whether the models and the data were suitable for cell type predictions. We used the same data and model to predict brain cell type proportions based on DNAm obtained from frontal brain tissue. Here, we were able to predict immune brain cell type composition in the independent testing dataset (**Figure S11**), suggesting that the combination of all DNAm sites in blood hold limited information on most brain cell types.

### 3.4 DNA methylation sites used in epigenomic cell deconvolution predicted brain cell types based on brain *and* blood data

The previous analyses focused either on single sites (EWAS) or a combination of sites without biological or statistical preselection of features (machine learning approaches). Next, we assessed whether a biologically relevant preselected combination of DNAm sites in peripheral blood could inform brain cell types. Of the 580 probes from the original brain cell reference panel selected to best predict cell proportions, 429 were present in all four frontal brain and blood datasets. We selected 48 DNAm sites that showed a significant correlation (Pearson’s r range: |0.198 - 0.537|) between frontal brain and peripheral blood in our curated superset (p<0.05) (**Table S5**). First, to confirm that these 48 DNAm sites in brain were still predictive of brain cell type proportions, we conducted a PCA for each of the seven cell types on the frontal brain DNAm data. To do so, we grouped the probes based on distinguishing power between cell types as indicated by the brain reference panel. For all these seven analyses, the first PC explained 98-100% of the variance for each cell type proportion, and they were significantly correlated with their corresponding brain cell type proportions (Pearson’s r: |0.571 - 0.969|, mean absolute correlation = 0.793, SD = 0.142). This indicated that using only 11.2% (48 out of 429) of probes in the brain cell type reference panel was still informative for brain cell type composition in the frontal brain (**Figure S12**). Then, we conducted the same analyses using blood DNAm and found that the first PC derived from the 48 blood-brain correlating probes in blood also explained variance of 98 to 100% in brain cell type proportions. This first PC was correlated with endothelial cells (Pearson’s r = 0.400, adjusted p-value = 1.78*10^-4^), microglia (Pearson’s r = 0.247, adjusted p-value = 0.0268), and stromal cells (Pearson’s r = 0.341, adjusted p-value = 1.40*10^-3^) (**Figure S12**). Given that these DNAm sites were chosen based on their strong single site DNAm correlations between blood and brain, it is perhaps not surprising that they performed well in predicting brain cell type proportions in the more complex PCA done here, However, we note that these specific CpGs only encompass less than 10% of the sites included in the deconvolution algorithm. Collectively, these relationships were in general agreement with our earlier findings of blood DNAm patterns being most predictive for immune-related cell types in the brain.

### 3.5 Correlations between blood immune cell types and brain cell type proportions were dependent on chronological age

Given the biological crosstalk between the peripheral immune system and brain, we next added an even more coarse, yet perhaps most physiologically relevant, layer to our suite of analyses - that being the direct comparison of predicted cell type proportions between blood and brain. With epigenomic deconvolution based on an extended blood cell type panel, we estimated twelve different immune cell types in blood (**Figure S14**) and then correlated these with our seven predicted brain cell type proportions. Despite many nominal mid-range correlations, after multiple test corrections, only NK cells were negatively correlated with stromal cells (Pearson’s r = 0.334, adjusted p-value = 0.044 (12 blood cell types x 7 brain cell types = 84 tests, **Figure S15**). However, since proportions of both cell types are relatively low, caution should be taken with the biological interpretation of this correlation.

Since the age of individuals included in the superset covered the whole lifespan and cell type proportions and their interactions change over the lifespan, we next investigated the effect of chronological age on the correlation between blood and brain cell type proportions (**Figure 5**). When residualizing cell type proportions on chronological age, the nominal correlations between cell types disappeared after multiple test correction (adjusted p-value > 0.05).

**Figure 5.**
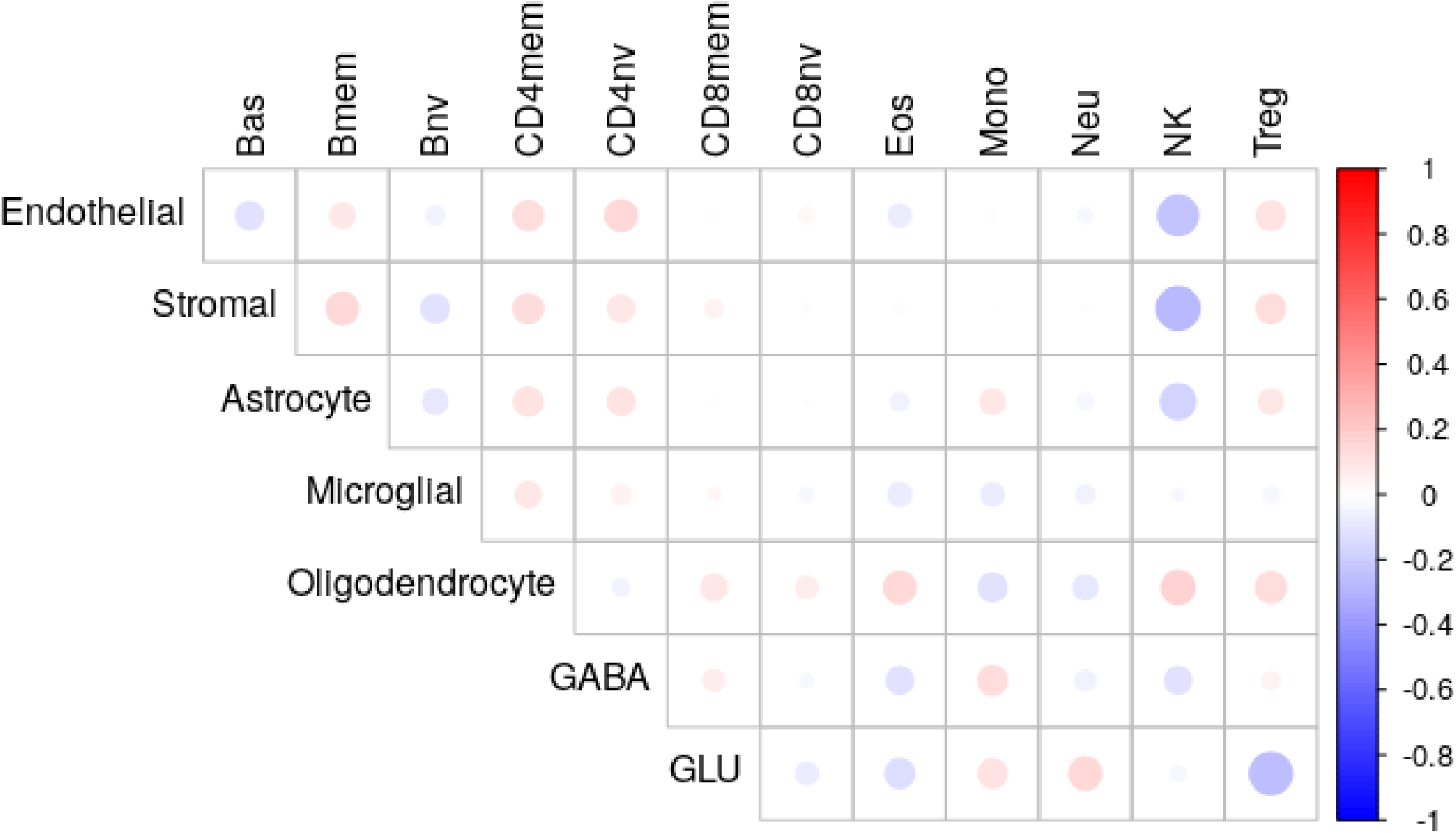
Blood and brain cell type proportions were not correlated with each other. Pearson’s correlations were calculated between brain cell type proportions (y-axis) and blood cell types (x-axis) both residualized on age from matched individuals, based on epigenomic deconvolution estimates. Size and colour of the circles indicate size of the Pearson’s correlation (bright red: Pearson’s r = 1, white: Pearson’s r=0, dark blue: Pearson’s r=-1).

### 3.6 Biological brain ageing was not correlated with brain or blood cell type proportions

The limited correlations between blood and brain cell type proportions were to some extent sensitive to age (**Figure 5**). However, age measures were based on chronological age, rather than biological aging which can be captured as EAA in the brain, which is associated with multiple brain-related diseases. We leveraged the cortical epigenetic clock to estimate epigenetic age followed by EAA (**Figure S16**). We assessed which cell type proportions could be driving cortical EAA. There was one potential outlier with a cortical EAA of 50, which was removed from subsequent analyses. However, cortical EAA was not correlated with any brain cell type (**Figure 6**).

**Figure 6.**
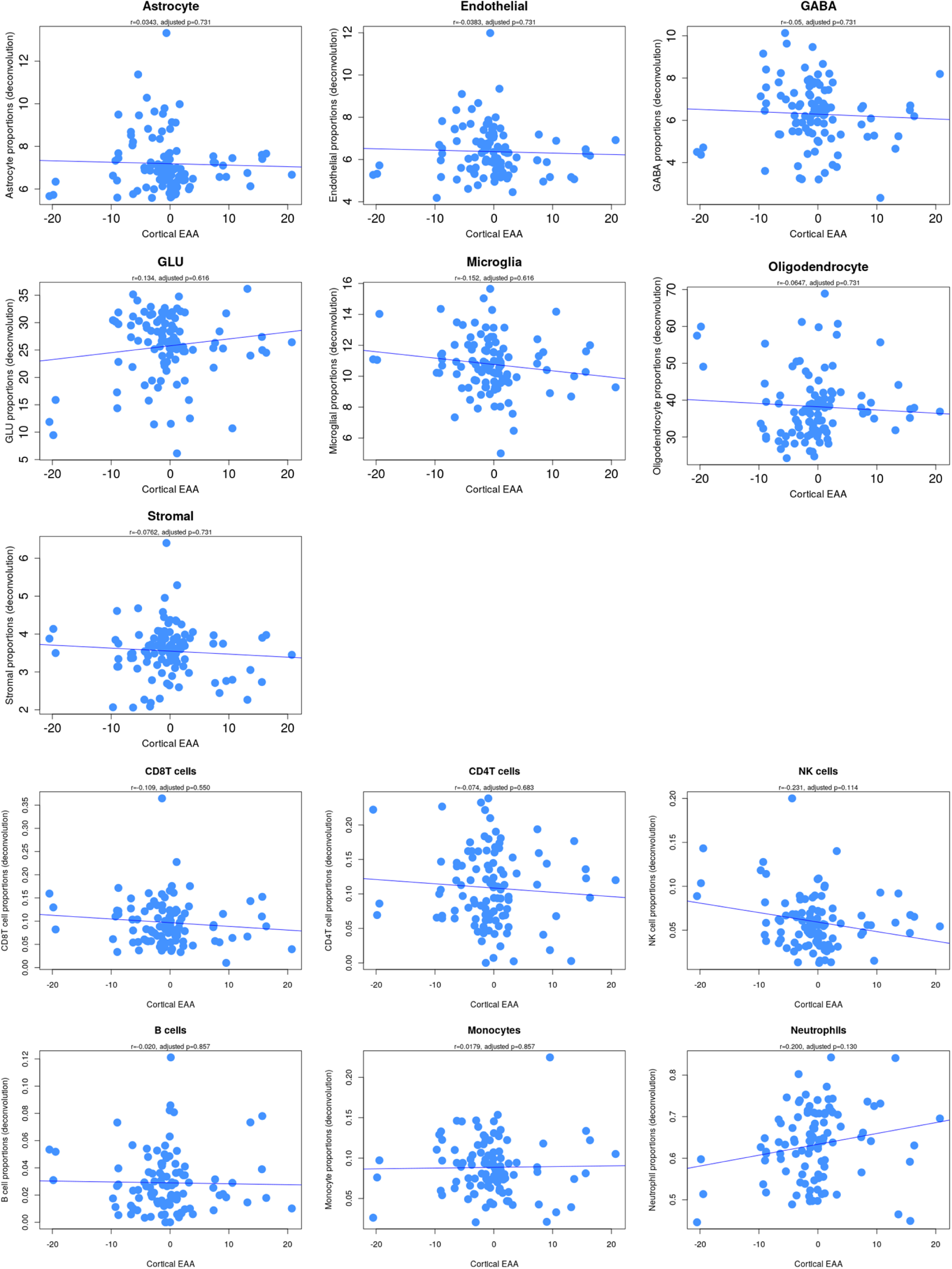
Biological ageing was not correlated with brain and blood cell type proportions. Brain and blood cell type proportions as estimated with epigenomic deconvolution are plotted on the y-axis. Cortical epigenetic age acceleration (EAA), defined as epigenetic age residualized on chronological age is plotted on the x-axis. Every dot represents a single sample in the combined dataset. The Pearson’s correlation (r) and p-value (p) are given for every brain cell type.

Previous research has shown that peripheral epigenetic signatures, including immune-related signatures, are associated with structural variation in brain and brain age acceleration (81, 82). We tested whether blood cell type proportions estimated based on peripheral epigenetic profiles could also serve as peripheral markers of brain ageing by assessing the association between cortical EAA and peripheral immune cell types in blood. Cortical EAA was also not significantly associated with blood cell type proportions (adjusted p-value < 0.05, **Figure 6**).

## 4. Discussion

In the current study, we investigated whether DNAm profiles in peripheral blood could be informative for deconvolution-estimated brain cell type proportions, given their implications in neurodevelopment, neurodegeneration, and psychiatric disorders. Using a comprehensive analytical strategy and utilizing bioinformatic predictors of cell type proportions, we investigated how DNAm in peripheral blood reflects cell type and epigenetic age measures in the brain. We first identified single DNAm sites in both brain and blood that were associated with brain cell type proportions, and then moved to multivariate models to see whether combining multiple DNAm sites hold more information over single sites. First, the number of brain-derived DNAm sites associated with predicted brain cell type proportions greatly outnumbered those derived from blood. Second, a multivariate predictor of brain cell type proportions created from blood DNAm sites showed the strongest predictive value for microglia. Related, at a coarser level, immune cell proportions from blood were correlated to corresponding immune-like cells from brain. Lastly, incorporating chronological and biological aging measures, we found that there was no association between EAA and any cell type. Collectively, our data suggested, specifically based on the PC-approach, that peripheral blood DNAm might be used to inform the proportion of immune-related brain cells.

We identified DNAm sites in frontal brain tissue that were associated with deconvolution-estimated brain cell type proportions, in similar amounts as have been discovered for EWAS in blood for isolated blood cell types (83–86). In our EWAS, we found that DNAm levels were consistently negatively associated with astrocytes, microglia, and stromal cell proportions. Associations between GABAergic neuron proportions and DNAm were mainly positive in the frontal brain. These findings agreed with previous reports showing that GABAergic neurons have higher global DNAm levels than glutamatergic neurons in the prefrontal cortex (87), and that glial cells (including astrocytes and microglia) have the lowest DNAm levels compared to neuronal cell populations in both the human (87, 88) and mouse brain (89). Furthermore, previous studies investigating blood DNAm in association with isolated blood cell types consistently showed that myeloid cells showed lower DNAm levels than lymphoid cells (83–86). Microglia also have a myeloid developmental origin. Next, we observed that considerable DNAm sites were associated with both oligodendrocytes and glutamatergic neurons. This overlap can best be explained by the fact that both cell type proportions are most common (90, 91), and negatively correlated with each other. In other words, even though the significant DNAm sites shared between oligodendrocytes and glutamatergic neurons are the same, these sites have effect sizes with opposite directions for each cell type.

In the previously discussed blood EWAS, the most abundant cell types (i.e., granulocytes and monocytes) also generally had the largest numbers of DNAm sites distinguishing them from the other cell types (83–86). We also identified that ∼10% of the DNAm sites that were associated with brain cell type proportions had significantly correlated DNAm levels across brain and blood. Previous studies show that 6-11% of all measured probes from either array (i.e., 450K or EPIC) exhibit peripheral blood-brain DNAm correlation (33–36). This indicates that DNAm sites in the frontal brain associated with deconvolution-estimated brain cell type proportions exhibited comparable levels of peripheral blood-brain correlation with what can be expected from the current array platforms available.

In peripheral blood, we identified a total of three statistically significant single DNAm sites associated with brain endothelial and stromal cell proportions, with small effect sizes (<0.04). These single DNAm sites also showed statistically significant peripheral blood-brain DNAm correlation. It is worth noting that the part of the HiBED deconvolution tool for two specific cell types, endothelial and stromal cells, was constructed based on cells isolated from infant cord peripheral blood (75). Because of the shared developmental origins of stromal and endothelial cells in blood and brain, the tissue differences can potentially drive a similar DNAm profile in the cell library and between peripheral and stromal cells in our combined dataset. Taken together, the DNAm associations highlight that some DNAm from both the frontal brain as well as peripheral blood were informative for deconvolution-estimated brain cell type proportions, potentially reflecting brain function, and possibly hinting at biological and physical interaction of systemic inflammation with the immune system in the brain.

We not only identified an association between peripheral DNAm profiles and immune brain cell type proportions, but we also identified associations between deconvolution-estimated peripheral immune cell proportions and different deconvolution-estimated brain cell types. We identified a nominally significant negative association between peripheral NK cell proportions and estimated brain stromal cell proportions. This finding should be interpreted with caution given that the association no longer holds up after multiple testing correction. Nevertheless, it raises a hypothesis worth pursuing: NK cells have the capacity of disrupt the blood-brain barrier (BBB) and to kill stromal cells lining the BBB in neuroinflammation (112). Replication in larger independent samples, followed by functional experiments are needed before any mechanistic conclusions can be drawn.

In our multi-levelled analysis, we used both single site and multivariate modelling to investigate whether DNAm sites in peripheral blood could predict brain cell type proportions. These approaches, especially based on PCs, revealed that peripheral blood DNAm sites were most predictive of immune cell type proportions in the brain, specifically microglia, and stromal cells. The fact that peripheral blood DNAm is informative for estimating immune brain cell type proportions is interesting as potential means of using peripheral blood DNAm to study the brain’s immune system, which is difficult to sample. Not only is brain tissue difficult to access, but its constituent cell types are also challenging to interrogate directly. For example, direct measurements with microscopy or flow cytometry are rarely performed. Using DNAm profiles derived from brain-tissue, we were able to estimate the cellular makeup in higher resolution and investigate its association with blood DNAm. However, it is important to reiterate that the cell type proportions are deconvolution-estimated and, despite the high reported accuracy (75, 76) can produce noisy and possibly erroneous data. That said, our findings may inform future studies using direct measurement approaches to target glial, endothelial, and stromal cells and the relationship with peripheral lymphocytes. The observed similarities between the two tissues might also indicate shared underlying mechanisms linking the brain and peripheral blood immune systems in the context of mental health. Consistent with this, the peripheral immune system is associated with different brain-related conditions such as schizophrenia (92), autism spectrum disorder (93), and AD (94) based on epidemiology, genetic correlation, and multi-omics approaches.

Different peripheral immune cell types, including the NK cells discussed above, have been found to be associated with epigenetic ageing measured with peripheral blood DNAm (72). Therefore, we investigated whether a similar association between epigenetic ageing and cell types, especially immune cells, also exists in the frontal brain. There was no association between epigenetic ageing of the brain and different deconvolution-estimated brain and blood cell types after multiple test correction, even though previous research showed that NK cells and neutrophils are negatively and positively associated with EAA and other measures of epigenetic age such PhenoAge and Pace of Ageing (72, 98, 99).

The results from the current study should be interpreted with their own limitations. First of all, we made use of publicly available databases, and all studies included different types of brain tissue from demographically different populations. For example, in the combined dataset we created, we included both postmortem and live brain tissue. It has been shown that postmortem and live brain tissue exhibit different gene expression patterns (100) and post-transcriptional RNA modifications (101), and it could thus be argued that epigenetic modifications, including DNAm might also be different between the two types of brain tissues. However, we did not observe substantial differences in cell type proportions between the individual cohorts, suggesting that the two types of tissues were similar enough for comparisons, especially because the normalization and batch correction steps of our preprocessing pipeline should have removed these effects. This is further supported by a previous study showing that the percentage of DNAm was stable up to 72 hours postmortem interval (96). Relatedly, not all matched blood and brain samples were collected at the same time point, with a median distance of two years between tissue collection from the same individual. For future data collection, timing of sample collection should be considered carefully. Second, we applied a machine learning algorithm to train a predictor of brain cell type proportions using peripheral DNAm profiles. We used three datasets with a total n=88 for training, and validated our findings in an independent dataset (n=16). The estimates obtained based on a sample size of N=104 distinct individuals with diverse demographics in the current study may be improved by an increase in sample size. The importance of statistical power is also underlined by our findings that the highest number of significantly associated DNAm sites were identified in oligodendrocytes, the most variable cell type in the frontal brain. Increases in sample size would also increase statistical power. Third, since our analysis was limited to frontal brain samples, and cell type proportions vary across brain regions (19, 103, 104), these findings may not generalize beyond the frontal brain. To draw robust conclusions about other brain regions, multiple matched brain regions and their individual correlation with blood should be further explored. Fourth, there was a variety in age range (e.g., only elderly individuals or a mix of adolescents and adults) across the sub-cohorts included in the current study. Our downstream data analyses were corrected for age, and therefore represent general effects across a 50-year age span. Thus, when considering research cohorts with a narrower age-range, it should be noted that correlations in one age group might not exist in another age group. Lastly, it is well-known that the commercial arrays do not distinguish between DNAm and DNA hydroxymethylation (DNAhm) (105). Given the importance of DNAhm in the brain (100), important brain cell type-associations could have been masked, as both epigenetic modifications show cell type-specific patterns in the brain (107). Furthermore, the difference between DNAm and DNAhm levels in both brain and blood would be important to take into account when investigating epigenetic correlations between tissues.

Based on the results presented in our study, we provide future directions for using DNAm as a tool to create cross-tissue biomarkers for unmeasured biological variables (108), such as brain cell type proportions. A larger sample size and the combination of both DNAm and DNAhm is recommended when aiming to develop such biomarkers. We also provided a short- list of DNAm sites which overlap on the Illumina 450K and EPICv1 arrays showing high peripheral blood-brain correlation *and* associations with brain cell type proportions (**Supplementary Table 5**). These DNAm sites can be used to identify the associations of brain cell type proportions involved in the phenotype of interest when performing association analyses in peripheral blood. Our study also showcased an example workflow to analyze diverse sets of cohorts. All in all, these results and recommendations open up the avenue for developing predictors of brain cell type proportions with larger sample size without the need to access brain tissue, allowing for more in-depth studies of biological pathways that underpin brain health and other brain-related phenotypes such as psychiatric/neurodevelopmental disorders and neurodegenerative diseases.

## Conclusions

In summary, through the lens of DNAm, the results presented here support previous studies finding an overlap between peripheral blood and the brain, specifically in the immune system. We showed that peripheral DNAm can be informative for immune brain cell type proportion estimates. Given the existing DNAm correlation between peripheral blood and brain, the usage of a stable cell identity measure, i.e., DNAm, in peripheral blood as biomarker of some brain cell type proportions could be plausible. Altogether, these results and recommendations could add to the interpretation of peripheral blood-derived DNAm-associations in the context of brain-related outcomes.

## Abbreviations

AD: Alzheimer’s disease
ADHD: Attention-deficit/hyperactivity disorder
BBB: Blood-brain-barrier
DNAm: DNA methylation
DNAhm: DNA hydroxymethylation
EAA: Epigenetic age acceleration
EWAS: Epigenome-wide association study
FDR: False Discovery Rate
HiBED: Hierarchical Brain Extended Deconvolution
IQR: Interquartile range
NK cells: Natural Killer cells
PCA: Principal Component analysis
PD: Parkinson’s disease
RF: Random Forest
RMSE: Root Mean Squared Error
SD: Standard deviation
VMR: Variable Methylation Region

## Declarations

### Ethics approval and consent to participate

All samples used in the current manuscript were publicly available (datasets). Each individual study has obtained ethics approval and consent from patients and/or family to participate. The following information is obtained from the original publications.

GSE214901: “The study protocol was approved by the Ethics Committee of the University of Fukui, Japan (Assurance no. 20200028), Yamaguchi University School of Medicine, Japan (Assurance no. 2020–202), and Sugita Genpaku Memorial Obama Municipal Hospital (Assurance no. 2–7). Moreover, this study was carried out in accordance with the Declaration of Helsinki and the Ethical Guidelines for Clinical Studies of the Ministry of Health, Labour and Welfare of Japan. All participants provided either written informed consent or both informed consent and assent.” (Nishitani, Translation Psychiatry, 2023).

GSE95049: “The Research Ethics Board at the Douglas Mental Health University Institute approved the project. Signed informed consent was obtained for each subject from next of kin.” (Farré, Epigenetics & Chromatin, 2015).

GSE111165: “This study was approved by the University of Iowa’s Human Subjects Research Institution Review Board. Written informed consent was obtained.” (Braun, Translation psychiatry, 2019).

GSE59685: “Ethical approval for the study was provided by the NHS South East London REC 3. Matched blood samples collected before death were available for a subset of individuals (Supplementary Tables 1 and 2) as part of the Alzheimer’s Research UK funded study “Biomarkers of AD Neurodegeneration”, with informed consent according to the Declaration of Helsinki (1991). For validation purposes STG and PFC tissue was obtained from 144 individuals archived in the Mount Sinai Alzheimer’s Disease and Schizophrenia Brain Bank (http://icahn.mssm.edu/research/labs/neuropathology-and-brain-banking) and EC, STG and PFC samples from an additional 62 individuals archived in the Thomas Willis Oxford Brain Collection (http://www.medsci.ox.ac.uk/optima/information-for-patients-and-the-public/the-thomas-willis-oxford-brain-collection).” (Lunnon, Nature Neuroscience, 2014).

## Availability of data and materials

The datasets analysed during the current study are available in the Gene Expression Omnibus repository, https://www.ncbi.nlm.nih.gov/geo/. We used the datasets with the following GEO Accession codes: GSE214901; GSE95049; GSE111165; GSE59685.

## Authors’ contributions

MM conceptualized and designed the study, performed analyses, visualized, interpret, wrote the manuscript, and acquired funding. MPF, EIND, and HRE supported analyses related to cell type proportions, machine learning approaches, and brain ageing, respectively. MPF, EIND, HRE, and MHC interpret results and revised the manuscript. MSK provided supervision and computational resources. All authors read and approved the final manuscript.

## Acknowledgements

We would like to thank Dr. Keegan Korthauer for her helpful insights on performing machine learning analyses. This research utilized the *FlowSorted.BloodExtended.EPIC* software packages developed at Dartmouth College, which are governed by the licensing terms provided by Dartmouth Technology Transfer.

## References

1. Global Health Data Exchange, Institute for Health Metrics and Evaluation, 2023

2. Chene G, Beiser A, Au R, Preis SR, Wolf PA, Dufouil C, et al. Gender and incidence of dementia in the Framingham Heart Study from mid-adult life. Alzheimers Dement. 2015;11(3):310–20.

3. Seshadri S, Wolf PA, Beiser A, Au R, McNulty K, White R, et al. Lifetime risk of dementia and Alzheimer’s disease. The impact of mortality on risk estimates in the Framingham Study. Neurology. 1997;49(6):1498–504.

4. Driver JA, Logroscino G, Gaziano JM, Kurth T. Incidence and remaining lifetime risk of Parkinson disease in advanced age. Neurology. 2009;72(5):432–8.

5. Zubenko GS, Zubenko WN, McPherson S, Spoor E, Marin DB, Farlow MR, et al. A collaborative study of the emergence and clinical features of the major depressive syndrome of Alzheimer’s disease. Am J Psychiatry. 2003;160(5):857–66.

6. Starkstein SE, Mizrahi R, Power BD. Depression in Alzheimer’s disease: phenomenology, clinical correlates and treatment. Int Rev Psychiatry. 2008;20(4):382–8.

7. Burchill E, Watson CJ, Fanshawe JB, Badenoch JB, Rengasamy E, Ghanem DA, et al. The impact of psychiatric comorbidity on Parkinson’s disease outcomes: a systematic review and meta-analysis. Lancet Reg Health Eur. 2024;39:100870.

8. McGrath JJ, Lim CCW, Plana-Ripoll O, Holtz Y, Agerbo E, Momen NC, et al. Comorbidity within mental disorders: a comprehensive analysis based on 145 990 survey respondents from 27 countries. Epidemiol Psychiatr Sci. 2020;29:e153.

9. Schmitt A, Malchow B, Hasan A, Falkai P. The impact of environmental factors in severe psychiatric disorders. Front Neurosci. 2014;8:19.

10. Tsankova N, Renthal W, Kumar A, Nestler EJ. Epigenetic regulation in psychiatric disorders. Nat Rev Neurosci. 2007;8(5):355–67.

11. Andreassen OA, Hindley GFL, Frei O, Smeland OB. New insights from the last decade of research in psychiatric genetics: discoveries, challenges and clinical implications. World Psychiatry. 2023;22(1):4–24.

12. Pihlstrom L, Wiethoff S, Houlden H. Genetics of neurodegenerative diseases: an overview. Handb Clin Neurol. 2017;145:309–23.

13. Cannon JR, Greenamyre JT. The role of environmental exposures in neurodegeneration and neurodegenerative diseases. Toxicol Sci. 2011;124(2):225–50.

14. Greenhalgh AD, David S, Bennett FC. Immune cell regulation of glia during CNS injury and disease. Nat Rev Neurosci. 2020;21(3):139–52.

15. Bradl M, Lassmann H. Oligodendrocytes: biology and pathology. Acta Neuropathol. 2010;119(1):37–53.

16. Yap CX, Vo DD, Heffel MG, Bhattacharya A, Wen C, Yang Y, et al. Brain cell-type shifts in Alzheimer’s disease, autism, and schizophrenia interrogated using methylomics and genetics. Sci Adv. 2024;10(21):eadn7655.

17. Maitra M, Mitsuhashi H, Rahimian R, Chawla A, Yang J, Fiori LM, et al. Cell type specific transcriptomic differences in depression show similar patterns between males and females but implicate distinct cell types and genes. Nat Commun. 2023;14(1):2912.

18. Saavedra J, Nascimento M, Liz MA, Cardoso I. Key brain cell interactions and contributions to the pathogenesis of Alzheimer’s disease. Front Cell Dev Biol. 2022;10:1036123.

19. Hannon E, Dempster EL, Davies JP, Chioza B, Blake GET, Burrage J, et al. Quantifying the proportion of different cell types in the human cortex using DNA methylation profiles. BMC Biol. 2024;22(1):17.

20. Pak V, Adewale Q, Bzdok D, Dadar M, Zeighami Y, Iturria-Medina Y. Distinctive whole-brain cell types predict tissue damage patterns in thirteen neurodegenerative conditions. Elife. 2024;12.

21. Stockmeier CA, Rajkowska G. Cellular abnormalities in depression: evidence from postmortem brain tissue. Dialogues Clin Neurosci. 2004;6(2):185–97.

22. Nestler EJ, Barrot M, DiLeone RJ, Eisch AJ, Gold SJ, Monteggia LM. Neurobiology of depression. Neuron. 2002;34(1):13–25.

23. Wang Q, Wang M, Choi I, Sarrafha L, Liang M, Ho L, et al. Molecular profiling of human substantia nigra identifies diverse neuron types associated with vulnerability in Parkinson’s disease. Sci Adv. 2024;10(2):eadi8287.

24. Skene NG, Bryois J, Bakken TE, Breen G, Crowley JJ, Gaspar HA, et al. Genetic identification of brain cell types underlying schizophrenia. Nat Genet. 2018;50(6):825–33.

25. Bryois J, Skene NG, Hansen TF, Kogelman LJA, Watson HJ, Liu Z, et al. Genetic identification of cell types underlying brain complex traits yields insights into the etiology of Parkinson’s disease. Nat Genet. 2020;52(5):482–93.

26. Price KM, Wigg KG, Nigam A, Feng Y, Blokland K, Wilkinson M, et al. Identification of brain cell types underlying genetic association with word reading and correlated traits. Mol Psychiatry. 2023;28(4):1719–30.

27. Meijer M, Franke B, Sandi C, Klein M. Epigenome-wide DNA methylation in externalizing behaviours: A review and combined analysis. Neurosci Biobehav Rev. 2023;145:104997.

28. Li E. Chromatin modification and epigenetic reprogramming in mammalian development. Nat Rev Genet. 2002;3(9):662–73.

29. Bird A. DNA methylation patterns and epigenetic memory. Genes Dev. 2002;16(1):6–21.

30. Feinberg AP, Irizarry RA, Fradin D, Aryee MJ, Murakami P, Aspelund T, et al. Personalized epigenomic signatures that are stable over time and covary with body mass index. Sci Transl Med. 2010;2(49):49ra67.

31. Pascual-Ahuir A, Fita-Torro J, Proft M. Capturing and Understanding the Dynamics and Heterogeneity of Gene Expression in the Living Cell. Int J Mol Sci. 2020;21(21).

32. Nigg JT. Considerations toward an epigenetic and common pathways theory of mental disorder. J Psychopathol Clin Sci. 2023;132(3):297–313.

33. Edgar RD, Jones MJ, Meaney MJ, Turecki G, Kobor MS. BECon: a tool for interpreting DNA methylation findings from blood in the context of brain. Transl Psychiatry. 2017;7(8):e1187.

34. Braun PR, Han S, Hing B, Nagahama Y, Gaul LN, Heinzman JT, et al. Genome-wide DNA methylation comparison between live human brain and peripheral tissues within individuals. Transl Psychiatry. 2019;9(1):47.

35. Nishitani S, Isozaki M, Yao A, Higashino Y, Yamauchi T, Kidoguchi M, et al. Cross-tissue correlations of genome-wide DNA methylation in Japanese live human brain and blood, saliva, and buccal epithelial tissues. Transl Psychiatry. 2023;13(1):72.

36. Hannon E, Lunnon K, Schalkwyk L, Mill J. Interindividual methylomic variation across blood, cortex, and cerebellum: implications for epigenetic studies of neurological and neuropsychiatric phenotypes. Epigenetics. 2015;10(11):1024–32.

37. Rovira P, Sanchez-Mora C, Pagerols M, Richarte V, Corrales M, Fadeuilhe C, et al. Epigenome-wide association study of attention-deficit/hyperactivity disorder in adults. Transl Psychiatry. 2020;10(1):199.

38. Hannon E, Dempster EL, Mansell G, Burrage J, Bass N, Bohlken MM, et al. DNA methylation meta-analysis reveals cellular alterations in psychosis and markers of treatment-resistant schizophrenia. Elife. 2021;10.

39. Aberg KA, McClay JL, Nerella S, Clark S, Kumar G, Chen W, et al. Methylome-wide association study of schizophrenia: identifying blood biomarker signatures of environmental insults. JAMA Psychiatry. 2014;71(3):255–64.

40. Cordova-Palomera A, Fatjo-Vilas M, Gasto C, Navarro V, Krebs MO, Fananas L. Genome-wide methylation study on depression: differential methylation and variable methylation in monozygotic twins. Transl Psychiatry. 2015;5(4):e557.

41. Dempster EL, Pidsley R, Schalkwyk LC, Owens S, Georgiades A, Kane F, et al. Disease-associated epigenetic changes in monozygotic twins discordant for schizophrenia and bipolar disorder. Hum Mol Genet. 2011;20(24):4786–96.

42. Roubroeks JAY, Smith AR, Smith RG, Pishva E, Ibrahim Z, Sattlecker M, et al. An epigenome-wide association study of Alzheimer’s disease blood highlights robust DNA hypermethylation in the HOXB6 gene. Neurobiol Aging. 2020;95:26–45.

43. Moore K, McKnight AJ, Craig D, O’Neill F. Epigenome-wide association study for Parkinson’s disease. Neuromolecular Med. 2014;16(4):845–55.

44. Dantzer R. Neuroimmune Interactions: From the Brain to the Immune System and Vice Versa. Physiol Rev. 2018;98(1):477–504.

45. Nudel R, Allesoe RL, Werge T, Thompson WK, Rasmussen S, Benros ME. An immunogenetic investigation of 30 autoimmune and autoinflammatory diseases and their links to psychiatric disorders in a nationwide sample. Immunology. 2023;168(4):622–39.

46. Berk M, Williams LJ, Jacka FN, O’Neil A, Pasco JA, Moylan S, et al. So depression is an inflammatory disease, but where does the inflammation come from? BMC Med. 2013;11:200.

47. Tylee DS, Sun J, Hess JL, Tahir MA, Sharma E, Malik R, et al. Genetic correlations among psychiatric and immune-related phenotypes based on genome-wide association data. Am J Med Genet B Neuropsychiatr Genet. 2018;177(7):641–57.

48. Collins JM, Keane JM, Deady C, Khashan AS, McCarthy FP, O’Keeffe GW, et al. Prenatal stress impacts foetal neurodevelopment: Temporal windows of gestational vulnerability. Neurosci Biobehav Rev. 2024;164:105793.

49. Jyonouchi H. Autism spectrum disorder and a possible role of anti-inflammatory treatments: experience in the pediatric allergy/immunology clinic. Front Psychiatry. 2024;15:1333717.

50. Enduru N, Fernandes BS, Bahrami S, Dai Y, Andreassen OA, Zhao Z. Genetic overlap between Alzheimer’s disease and immune-mediated diseases: an atlas of shared genetic determinants and biological convergence. Mol Psychiatry. 2024.

51. Ermakov EA, Melamud MM, Buneva VN, Ivanova SA. Immune System Abnormalities in Schizophrenia: An Integrative View and Translational Perspectives. Front Psychiatry. 2022;13:880568.

52. Saccaro LF, Crokaert J, Perroud N, Piguet C. Structural and functional MRI correlates of inflammation in bipolar disorder: A systematic review. J Affect Disord. 2023;325:83–92.

53. Vazquez-Gonzalez D, Carreon-Trujillo S, Alvarez-Arellano L, Abarca-Merlin DM, Dominguez-Lopez P, Salazar-Garcia M, et al. A Potential Role for Neuroinflammation in ADHD. Adv Exp Med Biol. 2023;1411:327–56.

54. Tansey MG, Wallings RL, Houser MC, Herrick MK, Keating CE, Joers V. Inflammation and immune dysfunction in Parkinson disease. Nat Rev Immunol. 2022;22(11):657–73.

55. Zenebe Y, Akele B, M WS, Necho M. Prevalence and determinants of depression among old age: a systematic review and meta-analysis. Ann Gen Psychiatry. 2021;20(1):55.

56. Schneider B, Prvulovic D, Oertel-Knochel V, Knochel C, Reinke B, Grexa M, et al. Biomarkers for major depression and its delineation from neurodegenerative disorders. Prog Neurobiol. 2011;95(4):703–17.

57. Weiskopf D, Weinberger B, Grubeck-Loebenstein B. The aging of the immune system. Transpl Int. 2009;22(11):1041–50.

58. Horvath S. DNA methylation age of human tissues and cell types. Genome Biol. 2013;14(10):R115.

59. Horvath S, Raj K. DNA methylation-based biomarkers and the epigenetic clock theory of ageing. Nat Rev Genet. 2018;19(6):371–84.

60. Shireby GL, Davies JP, Francis PT, Burrage J, Walker EM, Neilson GWA, et al. Recalibrating the epigenetic clock: implications for assessing biological age in the human cortex. Brain. 2020;143(12):3763–75.

61. Graves AJ, Danoff JS, Kim M, Brindley SR, Skyberg AM, Giamberardino SN, et al. Accelerated epigenetic age is associated with whole-brain functional connectivity and impaired cognitive performance in older adults. Sci Rep. 2024;14(1):9646.

62. Grodstein F, Lemos B, Yu L, Klein HU, Iatrou A, Buchman AS, et al. The association of epigenetic clocks in brain tissue with brain pathologies and common aging phenotypes. Neurobiol Dis. 2021;157:105428.

63. Levine ME, Lu AT, Bennett DA, Horvath S. Epigenetic age of the pre-frontal cortex is associated with neuritic plaques, amyloid load, and Alzheimer’s disease related cognitive functioning. Aging (Albany NY). 2015;7(12):1198–211.

64. Harvanek ZM, Boks MP, Vinkers CH, Higgins-Chen AT. The Cutting Edge of Epigenetic Clocks: In Search of Mechanisms Linking Aging and Mental Health. Biol Psychiatry. 2023;94(9):694–705.

65. Oblak L, van der Zaag J, Higgins-Chen AT, Levine ME, Boks MP. A systematic review of biological, social and environmental factors associated with epigenetic clock acceleration. Ageing Res Rev. 2021;69:101348.

66. Zhang ZZ, Moeckel C, Mustafa M, Pham H, Olson AE, Mehta D, et al. The association of epigenetic age acceleration and depressive and anxiety symptom severity among children recently exposed to substantiated maltreatment. J Psychiatr Res. 2023;165:7–13.

67. Han LKM, Aghajani M, Clark SL, Chan RF, Hattab MW, Shabalin AA, et al. Epigenetic Aging in Major Depressive Disorder. Am J Psychiatry. 2018;175(8):774–82.

68. Jeremian R, Malinowski A, Chaudhary Z, Srivastava A, Qian J, Zai C, et al. Epigenetic age dysregulation in individuals with bipolar disorder and schizophrenia. Psychiatry Res. 2022;315:114689.

69. Bourdon C, Etain B, Spano L, Belzeaux R, Leboyer M, Delahaye-Duriez A, et al. Accelerated aging in bipolar disorders: An exploratory study of six epigenetic clocks. Psychiatry Res. 2023;327:115373.

70. Salontaji K, Haftorn KL, Sanders F, Page CM, Walton E, Felix JF, et al. Gestational epigenetic age and ADHD symptoms in childhood: a prospective, multi-cohort study. Mol Psychiatry. 2024.

71. Postberg J, Schubert MT, Nin V, Wagner L, Piefke M. A perspective on epigenomic aging processes in the human brain and their plasticity in patients with mental disorders - a systematic review. Neurogenetics. 2024.

72. Jonkman TH, Dekkers KF, Slieker RC, Grant CD, Ikram MA, van Greevenbroek MMJ, et al. Functional genomics analysis identifies T and NK cell activation as a driver of epigenetic clock progression. Genome Biol. 2022;23(1):24.

73. Murthy M, Shireby G, Miki Y, Vire E, Lashley T, Warner TT, et al. Epigenetic age acceleration is associated with oligodendrocyte proportions in MSA and control brain tissue. Neuropathol Appl Neurobiol. 2023;49(1):e12872.

74. Gamo NJ, Arnsten AF. Molecular modulation of prefrontal cortex: rational development of treatments for psychiatric disorders. Behav Neurosci. 2011;125(3):282–96.

75. Zhang Z, Wiencke JK, Kelsey KT, Koestler DC, Molinaro AM, Pike SC, et al. Hierarchical deconvolution for extensive cell type resolution in the human brain using DNA methylation. Front Neurosci. 2023;17:1198243.

76. Salas LA, Zhang Z, Koestler DC, Butler RA, Hansen HM, Molinaro AM, et al. Enhanced cell deconvolution of peripheral blood using DNA methylation for high-resolution immune profiling. Nat Commun. 2022;13(1):761.

77. Houseman EA, Accomando WP, Koestler DC, Christensen BC, Marsit CJ, Nelson HH, et al. DNA methylation arrays as surrogate measures of cell mixture distribution. BMC Bioinformatics. 2012;13:86.

78. Willer CJ, Li Y, Abecasis GR. METAL: fast and efficient meta-analysis of genomewide association scans. Bioinformatics. 2010;26(17):2190–1.

79. Navarro-Delgado E, Czamara D, Edwards K, Fu MP, Merill SM, Konwar C, MacIsaac JL, Lin DTS, Mandhane P, Simons E, Subbarao P, Moraes TJ, Lahti J, Miller GR, Binder EB, Räikkönen K, Turvey SE, Korthauer K, Kobor MS, RAMEN: Regional Association of Methylome variability with the Exposome and geNome.bioRxiv 2025.05.08.652964;

80. Wright MN ZA. ranger: A Fast Implementation of Random Forests for High Dimensional Data in C++ and R. Journal of Statistical Software. 2017;77(1):1–17.

81. Freytag V, Carrillo-Roa T, Milnik A, Samann PG, Vukojevic V, Coynel D, et al. A peripheral epigenetic signature of immune system genes is linked to neocortical thickness and memory. Nat Commun. 2017;8:15193.

82. Sant’Anna Barbosa Ferreira P, van Dongen J, den Braber A, Boomsma DI, de Geus EJC, van ’t Ent D. Epigenetic age acceleration in peripheral blood correlates with brain-MRI age acceleration. Brain. 2025.

83. Roy R, Ramamoorthy S, Shapiro BD, Kaileh M, Hernandez D, Sarantopoulou D, Arepalli S, Boller S, Singh A, Bektas A, Kim J, Moore AZ, Tanaka T, McKelvey J, Zukley L, Nguyen C, Wallace T, Dunn C, Wersto R, Wood W, Piao Y, Becker KG, Coletta C, De S, Sen JM, Battle A, Weng NP, Grosschedl R, Ferrucci L, Sen R. DNA methylation signatures reveal that distinct combinations of transcription factors specify human immune cell epigenetic identity. Immunity. 2021 Nov 9;54(11):2465–2480.e5.

84. Bergstedt J, Azzou SAK, Tsuo K, Jaquaniello A, Urrutia A, Rotival M, Lin DTS, MacIsaac JL, Kobor MS, Albert ML, Duffy D, Patin E, Quintana-Murci L; Milieu Intérieur Consortium. The immune factors driving DNA methylation variation in human blood. Nat Commun. 2022 Oct 6;13(1):5895.

85. Reinius LE, Acevedo N, Joerink M, Pershagen G, Dahlén SE, Greco D, Söderhäll C, Scheynius A, Kere J. Differential DNA methylation in purified human blood cells: implications for cell lineage and studies on disease susceptibility. PLoS One. 2012;7(7):e41361.

86. Bakulski KM, Feinberg JI, Andrews SV, Yang J, Brown S, L McKenney S, Witter F, Walston J, Feinberg AP, Fallin MD. DNA methylation of cord blood cell types: Applications for mixed cell birth studies. Epigenetics. 2016 May 3;11(5):354–62.

87. Kozlenkov A, Wang M, Roussos P, Rudchenko S, Barbu M, Bibikova M, et al. Substantial DNA methylation differences between two major neuronal subtypes in human brain. Nucleic Acids Res. 2016;44(6):2593–612.

88. Tian W, Zhou J, Bartlett A, Zeng Q, Liu H, Castanon RG, et al. Single-cell DNA methylation and 3D genome architecture in the human brain. Science. 2023;382(6667):eadf5357.

89. Liu H, Zhou J, Tian W, Luo C, Bartlett A, Aldridge A, et al. DNA methylation atlas of the mouse brain at single-cell resolution. Nature. 2021;598(7879):120–8.

90. Hornung JP, De Tribolet N. Distribution of GABA-containing neurons in human frontal cortex: a quantitative immunocytochemical study. Anat Embryol (Berl). 1994 Feb;189(2):139–45.

91. Pelvig DP, Pakkenberg H, Stark AK, Pakkenberg B. Neocortical glial cell numbers in human brains. Neurobiol Aging. 2008 Nov;29(11):1754–62.

92. van Mierlo HC, Schot A, Boks MPM, de Witte LD. The association between schizophrenia and the immune system: Review of the evidence from unbiased ’omic-studies’. Schizophr Res. 2020;217:114–23.

93. Robinson-Agramonte MLA, Noris Garcia E, Fraga Guerra J, Vega Hurtado Y, Antonucci N, Semprun-Hernandez N, et al. Immune Dysregulation in Autism Spectrum Disorder: What Do We Know about It? Int J Mol Sci. 2022;23(6).

94. Bettcher BM, Tansey MG, Dorothee G, Heneka MT. Peripheral and central immune system crosstalk in Alzheimer disease - a research prospectus. Nat Rev Neurol. 2021;17(11):689–701.

95. Varatharaj A, Galea I. The blood-brain barrier in systemic inflammation. Brain Behav Immun. 2017;60:1–12.

96. Matsuoka RL, Buck LD, Vajrala KP, Quick RE, Card OA. Historical and current perspectives on blood endothelial cell heterogeneity in the brain. Cell Mol Life Sci. 2022;79(7):372.

97. Li Z, Li M, Shi SX, Yao N, Cheng X, Guo A, et al. Brain transforms natural killer cells that exacerbate brain edema after intracerebral hemorrhage. J Exp Med. 2020;217(12).

98. Zhang Z, Reynolds SR, Stolrow HG, Chen JQ, Christensen BC, Salas LA. Deciphering the role of immune cell composition in epigenetic age acceleration: Insights from cell-type deconvolution applied to human blood epigenetic clocks. Aging Cell. 2024;23(3):e14071.

99. Sun M, Yang H, Hu Y, Fan J, Duan M, Ruan J, et al. Differential white blood cell count and epigenetic clocks: a bidirectional Mendelian randomization study. Clin Epigenetics. 2024;16(1):118.

100. Collado-Torres L, Klei L, Liu C, Kleinman JE, Hyde TM, Geschwind DH, et al. Comparison of gene expression in living and postmortem human brain. medRxiv. 2023.

101. Rodriguez de Los Santos M, Kopell BH, Buxbaum Grice A, Ganesh G, Yang A, Amini P, et al. Divergent landscapes of A-to-I editing in postmortem and living human brain. Nat Commun. 2024;15(1):5366.

102. Rhein M, Hagemeier L, Klintschar M, Muschler M, Bleich S, Frieling H. DNA methylation results depend on DNA integrity-role of post mortem interval. Front Genet. 2015;6:182.

103. Siletti K, Hodge R, Mossi Albiach A, Lee KW, Ding SL, Hu L, et al. Transcriptomic diversity of cell types across the adult human brain. Science. 2023;382(6667):eadd7046.

104. Langlieb J, Sachdev NS, Balderrama KS, Nadaf NM, Raj M, Murray E, Webber JT, Vanderburg C, Gazestani V, Tward D, Mezias C, Li X, Cable DM, Norton T, Mitra P, Chen F, Macosko EZ. The cell type composition of the adult mouse brain revealed by single cell and spatial genomics. bioRxiv [Preprint]. 2023 Mar 13:2023.03.06.531307.

105. Huang Y, Pastor WA, Shen Y, Tahiliani M, Liu DR, Rao A. The behaviour of 5- hydroxymethylcytosine in bisulfite sequencing. PLoS One. 2010;5(1):e8888.

106. Kinney SM, Chin HG, Vaisvila R, Bitinaite J, Zheng Y, Esteve PO, et al. Tissue-specific distribution and dynamic changes of 5-hydroxymethylcytosine in mammalian genomes. J Biol Chem. 2011;286(28):24685–93.

107. Tooley KB, Chucair-Elliott AJ, Ocanas SR, Machalinski AH, Pham KD, Hoolehan W, et al. Differential usage of DNA modifications in neurons, astrocytes, and microglia. Epigenetics Chromatin. 2023;16(1):45.

108. Schmunk LJ, Call TP, McCartney DL, Javaid H, Hastings WJ, Jovicevic V, Kojadinović D, Tomkinson N, Zlamalova E, McGee KC, Sullivan J, Campbell A, McIntosh AM, Óvári V, Wishart K, Behrens CE, Stone E, Gavrilov M, Thompson R; Hurdle bio-infrastructure team; Jackson T, Lord JM, Stubbs TM, Marioni RE, Martin-Herranz DE. A novel framework to build saliva-based DNA methylation biomarkers: Quantifying systemic chronic inflammation as a case study. Aging Cell. 2025 Apr;24(4):e14444.

109. Fortin, J. P., Triche, T. J., Jr, & Hansen, K. D. (2017). Preprocessing, normalization and integration of the Illumina HumanMethylationEPIC array with minfi. Bioinformatics (Oxford, England), 33(4), 558–560. 10.1093/bioinformatics/btw691

110. Nevalainen, T., Autio, A., & Hurme, M. (2022). Composition of the infiltrating immune cells in the brain of healthy individuals: effect of aging. Immunity & ageing: I & A, 19(1), 45. 10.1186/s12979-022-00302-y

111. Netzahualcoyotzi, C., Santillán-Cigales, J. J., Adalid-Peralta, L. V., & Velasco, I. (2024). Infiltration of immune cells to the brain and its relation to the pathogenesis of Alzheimer’s and Parkinson’s diseases. Journal of neurochemistry, 168(9), 2316–2334. 10.1111/jnc.16106

112. Ning Z, Liu Y, Guo D, et al. (2023). Natural killer cells in the central nervous system. Cell Commun Signal 21: 341

