## Supplementary for "Peripheral blood DNA methylation partially reports on immune cell type composition in the frontal brain"

### Supplementary material

**Supplementary methods**

*DNA methylation*

For dataset 1 specifically, iDATs from the 450K and EPIC array were combined in a virtual 450K array through the *combineArrays* function from the same *minfi* package v1.44.0 [1]. Beta values were extracted from the RGset. DNAm quality control (QC) was performed in each dataset individually. Samples with a median methylated (M) and unmethylated (U) intensity below $\log_2(10.5)$ were removed, as well as samples with a bisulfite conversion rate < 0.8, and samples of which DNAm-based predicted sex did not match reported sex, assessed with the *GetSex* function in the *minfi* package. Correlation between 59 common single nucleotide polymorphisms (SNPs) was used to identify duplicate samples, and duplicated samples were removed. Samples with > 1% of probes with a detection p-value > 0.01 were removed, as well as samples with > 1% of probes with a bead count < 3. Probes located on the X- and Y-chromosomes were removed, as well as probes known to cross-hybridize on either the 450K and EPIC array or to contain SNPs with a minor allele frequency > 0.1 in the general population as defined by the 1K Genome project [2, 3]. Then, all probes with a bead count < 3 in > 1% samples, or > 1% of samples with a detection p-value > 0.05 were removed. For dataset 3, M and U intensity measures and detection p-values were available for processing. Samples with a median M and U intensity below $\log_2(10.5)$ were removed, as well as samples with predicted sex-mismatch and duplicated samples. Probe removal in dataset 3 was performed similarly as in datasets 1, 2, and 4. After sample and probe removal in the four different datasets, for each tissue type, only the overlapping probes in all four datasets were retained. Dasen normalization was performed on the superset for each tissue type [4], followed by *ComBat* v3.46.0 adjusting for slide and array platform (i.e., 450K and EPIC array) [5].

*Multivariate brain cell type proportion predictors based on blood DNAm data*

For ridge regression, the dependent outcomes (i.e., a specific brain cell type proportion) were centred, and a generalized linear model was fit on the peripheral blood beta values, with an alpha of 0 using the *glmnet* R package [6, 7]. Regularization hyperparameters to predict brain cell type proportions were tuned based on the 5-fold cross-validated models, choosing the lambda with the smallest mean squared error. All estimated cell type proportion with a negative value were constrained to 0. The same strategy was applied to the Ridge regression models based on brain DNAm sites.

VMRs can help to reduce the dimensionality of the data [8]. Within one VMR, the probe with the highest pairwise correlation with the other DNAm sites in the VMR was selected for Ridge regression. When a VMR was constituted by two probes, a random probe was selected. Following, a Random Forest (RF) model was built with the same probe selection criteria. Dependent outcomes were centred in the same way as with Ridge regression.

*Epigenetic ageing in the brain*

To calculate epigenetic age with the Cortical Clock [9], missing probes were imputed based on reference betas of the average DNAm levels across 700 control cortical samples [10]. Then, epigenetic brain age was calculated as the weighted sum of the coefficients for each probe plus the intercept, as described previously [9]. The logarithmic relationship observed in age 0-20 years was accounted for by anti-transforming the results [11]. EAA was calculated by residualizing predicted epigenetic brain age on chronological age. To identify the cell type-independent epigenetic aging signal of the frontal cortex, intrinsic brain EAA was calculated by residualizing both chronological age and all estimated brain cell type proportions on the predicted brain age.

### Supplementary results

*Table S1 Demographics of individuals in the sub-cohorts and superset.*

| | GSE | DNA methylation platform | Data source | Sample type | N | Age blood (mean (SD;range)) | Age brain (mean (SD;range)) | Male (%) | Source paper |
|---|---|---|---|---|---|---|---|---|---|
| Dataset 1 | GSE214901 | 450K and EPIC | University of Iowa Hospitals and Clinics, United States | Live brain tissue samples from patients with medically intractable epilepsy undergoing brain resection | 8 | 27.75 years (16.72; 5-60) | 27.75 years (16.72; 5-60) | 63% | Braun, 2019, Translation Psychiatry |
| Dataset 2 | GSE95049 | EPIC | University of Fukui Hospital, Yamaguchi University Hospital and Sugita Genpaku Memorial Obama Municipal Hospital, Japan | Live brain tissue samples from patients undergoing neurosurgery for their clinical purposes, including intractable epilepsy, meningioma, and cerebrovascular diseases | 6 | 65.33 (5.54; 59-73) | 65.33 (5.54; 59-73) | 50% | Nishitani, 2019, Translational Psychiatry |
| Dataset 3 | GSE59685 | 450K | MRC London Neurodegenerative Disease Brain Bank, United Kingdom | Post-mortem brain tissue dissected at autopsy | 74 | 83.32 (6.54; 70-99) | 87.15 (6.67; 71-105) | 35% | Hannon, 2015, epigenetics |
| Dataset 4 | GSE95049 | 450K | Douglas-Bell Canada Brain Bank, Canada | Post-mortem brain tissue dissected at autopsy | 16 | 57.50 (24.29; 15-87) | 57.50 (24.29; 15-87) | 75% | Farré, 2015, Epigenetics and Chromatin |
| Superset | | Virtual 450K | | | 104 | 74.04 (20.30; 5-99) | 76.76 (21.62; 5-105) | 44% | |

*Table S2 Number of surrogate variables included in the epigenome-wide association models for each cell type in each cohort*

| | Astrocyte | | Endothelial | | GABA | | GLU | | Microglia | | Oligodendrocyte | | Stromal | |
|---|---|---|---|---|---|---|---|---|---|---|---|---|---|---|
| | Brain | Blood | Brain | Blood | Brain | Blood | Brain | Blood | Brain | Blood | Brain | Blood | Brain | Blood |
| **Dataset 1** | 2 | 1 | 1 | 1 | 1 | 1 | 1 | 2 | 1 | 1 | 1 | 1 | 2 | 1 |
| **Dataset 2** | 1 | 1 | 1 | 1 | 0 | 0 | 1 | 1 | 1 | 1 | 0 | 0 | 1 | 0 |
| **Dataset 3** | 8 | 18 | 10 | 18 | 9 | 18 | 11 | 18 | 9 | 19 | 13 | 18 | 7 | 19 |
| **Dataset 4** | 1 | 4 | 1 | 4 | 1 | 3 | 4 | 3 | 1 | 4 | 5 | 4 | 2 | 4 |

Table S4 (Most) significant DNAm-associations in blood per brain cell type

| Cell type | CpG site | Gene | Z-score | P-value | Direction of effect | FDR | In deconvolution panel? | Blood-brain correlation |
|---|---|---|---|---|---|---|---|---|
| Astrocytes | cg13321164 | *GDF2* | 5.015 | $5.31*10^{-7}$ | ++++ | 0.195 | No | r=0.167, p=0.119 |
| Endothelial | cg02070873 | *ALOX5* | 5.672 | $1.41*10^{-8}$ | ++++ | 0.0362 | No | r=0.398, $p=1.21*10^{-4}$ |
| | cg23665824 | *CTNN2A; LRRTM1* | 5.518 | $3.44*10^{-8}$ | ++++ | 0.0441 | No | r=0.0334, p=0.757 |
| GABA | cg08099136 | *PSMB8* | -4.599 | $4.24*10^{-6}$ | ---- | 1 | No | r=0.108, p=0.314 |
| GLU | cg19613828 | *SASH1* | 4.978 | $6.41*10^{-6}$ | ++++ | 0.235 | No | r=0.0741, p=0.492 |
| Microglia | cg00580497 | Intergenic | 4.577 | $4.71*10^{-6}$ | +-++ | 1 | No | r=0.116, p=0.280 |
| Oligodendrocyte | cg22474886 | Intergenic | 4.948 | $7.49*10^{-7}$ | ++++ | 0.274 | No | r=-0.160, p=0.136 |
| Stromal | cg07078747 | *ALG10* | -6.016 | $1.79*10^{-9}$ | -+-- | 0.0046 | No | r=0.900, $p=1.06*10^{-32}$ |

Table S5 DNAm sites with blood-frontal cortex correlation present in the brain cell type deconvolution reference panel

| CpG site | Cell type reference panel | Blood-frontal cortex correlation (Pearson's r) | Blood-frontal cortex correlation (p-value) |
|---|---|---|---|
| cg00324161 | Astrocyte | 0.276 | $4.61*10^{-3}$ |
| cg00620628 | Glial; Neuronal | 0.291 | $2.77*10^{-3}$ |
| cg00872435 | Stromal | 0.208 | $3.40*10^{-2}$ |
| cg00968638 | Neuronal | 0.220 | $2.49*10^{-2}$ |
| cg01199952 | Oligodendrocyte | 0.286 | $3.29*10^{-3}$ |
| cg01604994 | Oligodendrocyte | -0.267 | $6.20*10^{-3}$ |
| cg01704924 | Stromal | 0.236 | $1.57*10^{-2}$ |
| cg02381853 | Astrocyte; Neuronal; Oligodendrocyte | -0.221 | $2.41*10^{-2}$ |
| cg02873421 | Endothelial and Stromal; Microglial | -0.336 | $4.81*10^{-4}$ |
| cg03312532 | Endothelial | 0.207 | $3.46*10^{-2}$ |
| cg03370270 | Neuronal | -0.265 | $6.65*10^{-3}$ |
| cg04979811 | GABA | 0.222 | $2.33*10^{-2}$ |
| cg05254095 | Oligodendrocyte | 0.203 | $3.91*10^{-2}$ |
| cg05268155 | Endothelial | 0.210 | $3.21*10^{-2}$ |
| cg05347108 | Astrocyte; Endothelial and Stromal; Microglial | -0.365 | $1.40*10^{-4}$ |
| cg05527091 | GLU | -0.235 | $1.65*10^{-2}$ |
| cg06023661 | Astrocyte | 0.226 | $2.11*10^{-2}$ |
| cg06710464 | GLU | 0.245 | $1.21*10^{-2}$ |
| cg06934394 | Glial, Neuronal | 0.197 | $4.55*10^{-2}$ |
| cg07106927 | Astrocyte | 0.205 | $3.72*10^{-2}$ |
| cg07730301 | Glial, Stromal | -0.241 | $1.38*10^{-2}$ |
| cg07912020 | Astrocyte | 0.225 | $2.16*10^{-2}$ |
| cg08331427 | Neuronal | 0.676 | $3.24*10^{-5}$ |
| cg08841511 | Neuronal | 0.236 | $1.60*10^{-2}$ |
| cg09216650 | Astrocyte | 0.228 | $2.01*10^{-2}$ |
| cg09404376 | Astrocyte | 0.214 | $2.93*10^{-2}$ |
| cg10432083 | Glial | 0.239 | $1.46*10^{-2}$ |
| cg10436631 | Astrocyte | 0.577 | $1.45*10^{-10}$ |
| cg11165158 | Endothelial | 0.208 | $3.39*10^{-2}$ |
| cg12634591 | Endothelial | 0.260 | $7.62*10^{-3}$ |
| cg13189759 | GLU | 0.223 | $2.27*10^{-2}$ |
| cg13400443 | Endothelial | 0.202 | $3.97*10^{-2}$ |
| cg14030346 | Endothelial | 0.214 | $2.94*10^{-2}$ |
| cg15687622 | GLU | 0.222 | $2.35*10^{-2}$ |
| cg15850851 | Neuronal | 0.277 | $4.35*10^{-3}$ |
| cg15894722 | Stromal | 0.381 | $6.56*10^{-5}$ |
| cg16533359 | Glial | 0.233 | $1.71*10^{-2}$ |
| cg17153363 | GABA | 0.305 | $1.67*10^{-3}$ |
| cg17714703 | Endothelial | 0.447 | $1.96*10^{-6}$ |
| cg17818471 | Endothelial and Stromal | 0.414 | $1.23*10^{-5}$ |
| cg18011078 | Glial | 0.221 | $2.40*10^{-2}$ |
| cg18215449 | Neuronal | -0.198 | $4.34*10^{-2}$ |
| cg20445094 | Neuronal | -0.223 | $2.31*10^{-2}$ |
| cg21270243 | Endothelial and Stromal | 0.290 | $2.83*10^{-3}$ |
| cg23389651 | Neuronal | -0.223 | $2.31*10^{-2}$ |
| cg26175287 | Neuronal; Oligodendrocyte | 0.404 | $2.09*10^{-5}$ |
| cg26644853 | Microglial | 0.314 | $1.17*10^{-3}$ |

| | | | |
|---|---|---|---|
| cg27626141 | Neuronal; Oligodendrocyte | 0.537 | $4.30*10^{-9}$ |

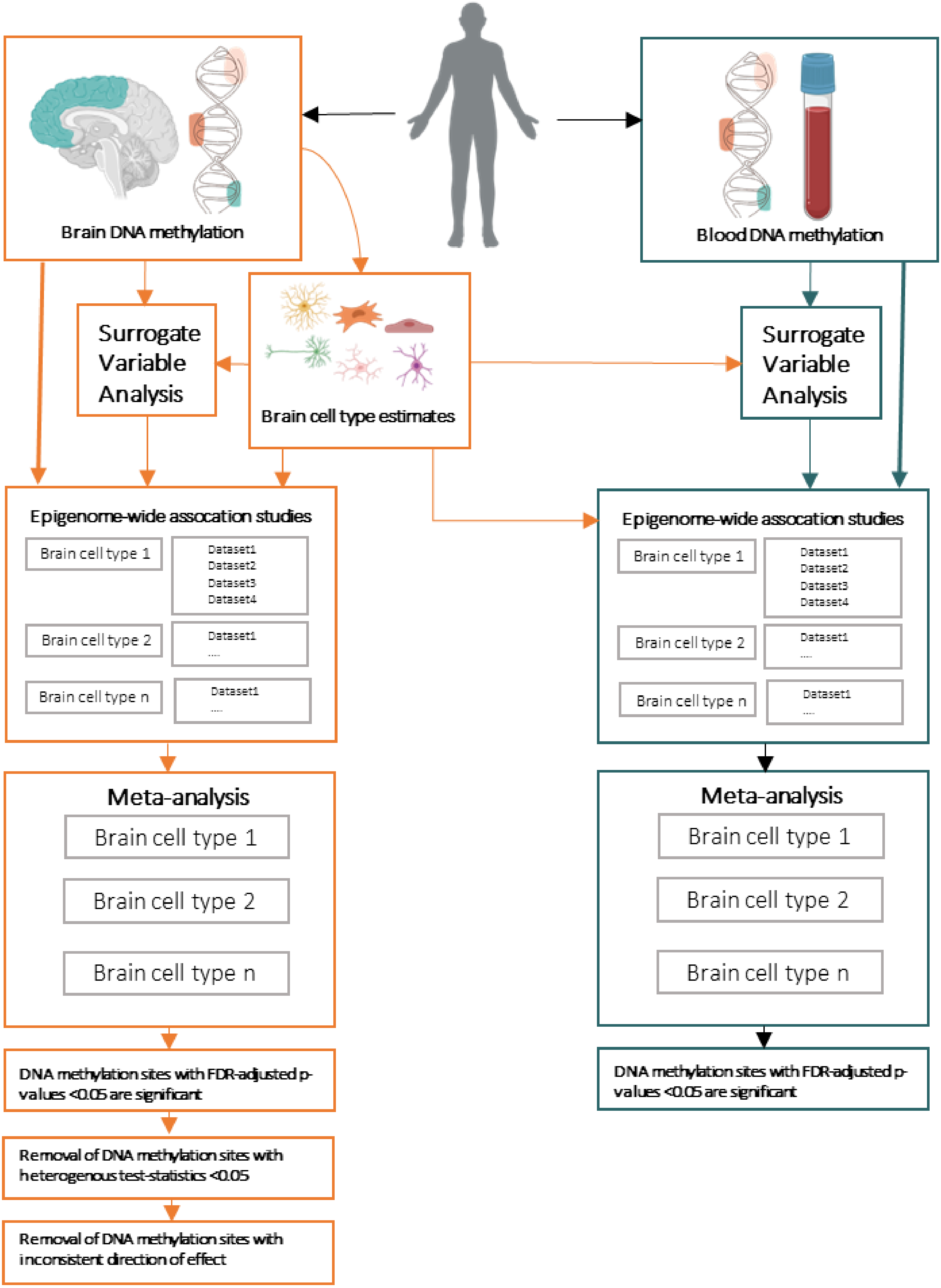
Brain DNA methylation
Blood DNA methylation
Surrogate Variable Analysis
Brain cell type estimates
Surrogate Variable Analysis
Epigenome-wide assocation studies
Brain cell type 1
Dataset1
Dataset2
Dataset3
Dataset4
Brain cell type 2
Dataset1
....
Brain cell type n
Dataset1
....
Epigenome-wide assocation studies
Brain cell type 1
Dataset1
Dataset2
Dataset3
Dataset4
Brain cell type 2
Dataset1
....
Brain cell type n
Dataset1
....
Meta-analysis
Brain cell type 1
Brain cell type 2
Brain cell type n
Meta-analysis
Brain cell type 1
Brain cell type 2
Brain cell type n
DNA methylation sites with FDR-adjusted p-values <0.05 are significant
DNA methylation sites with FDR-adjusted p-values <0.05 are significant
Removal of DNA methylation sites with heterogenous test-statistics <0.05
Removal of DNA methylation sites with inconsistent direction of effect

***Figure S1 Flowchart of epigenome-wide association meta-analysis for brain cell types in brain and blood.*** *Brain and blood DNA methylation was used to estimate brain- and blood cell type proportions. Surrogate variable analysis (SVA) was performed to determine the amount of surrogate variables to include in the statistical models for each epigenome-wide association study (EWAS;* ***Table S2****). An EWAS was performed for all brain cell types, in each cohort separately, both in brain tissue and peripheral blood. Results from the individual EWAS were meta-analysed per cell type per tissue. DNA methylation sites with an FDR-adjusted p-value <0.05 were considered staticsally significant. For the meta-analyses in brain tissue, DNA methylation sites with heterogeneous test-statistics <0.05 were removed, as well as DNA methylation sites with inconsistent direction of effects.*

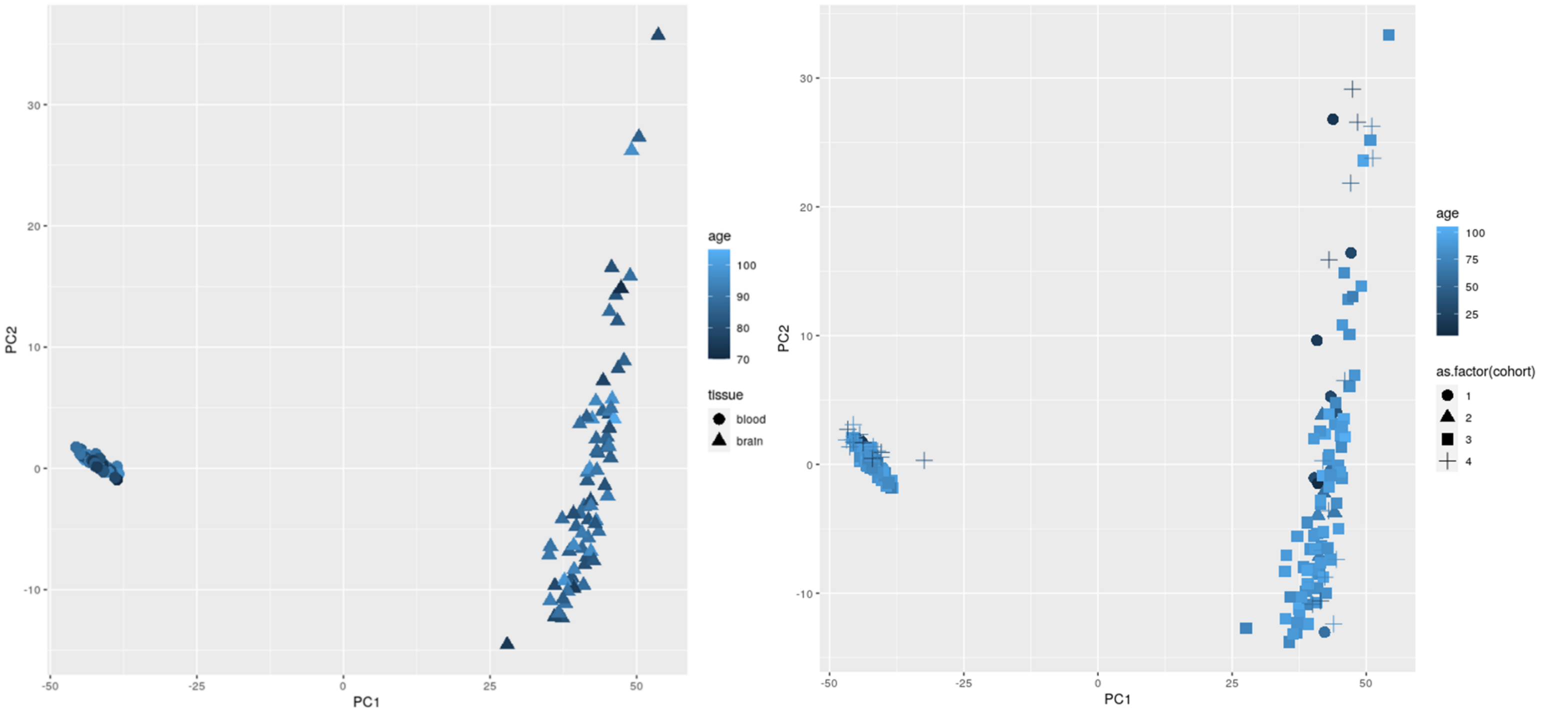


***Figure S2 Tissue differences explained most variance in DNA methylation levels in all cohorts.*** *Left panel: Principal component analysis (PCA) was performed on the DNA methylation beta values of blood (circle) and brain (triangle) in cohort 3 (i.e., cohort with up to 16-year time gap between data collection points). Colour gradients reflect chronological age of tissue collection. Right panel: PCA was performed on the DNA methylation beta values of all four cohorts (cohort indicated by signs), and colour gradients reflect chronological age at tissue collection.*

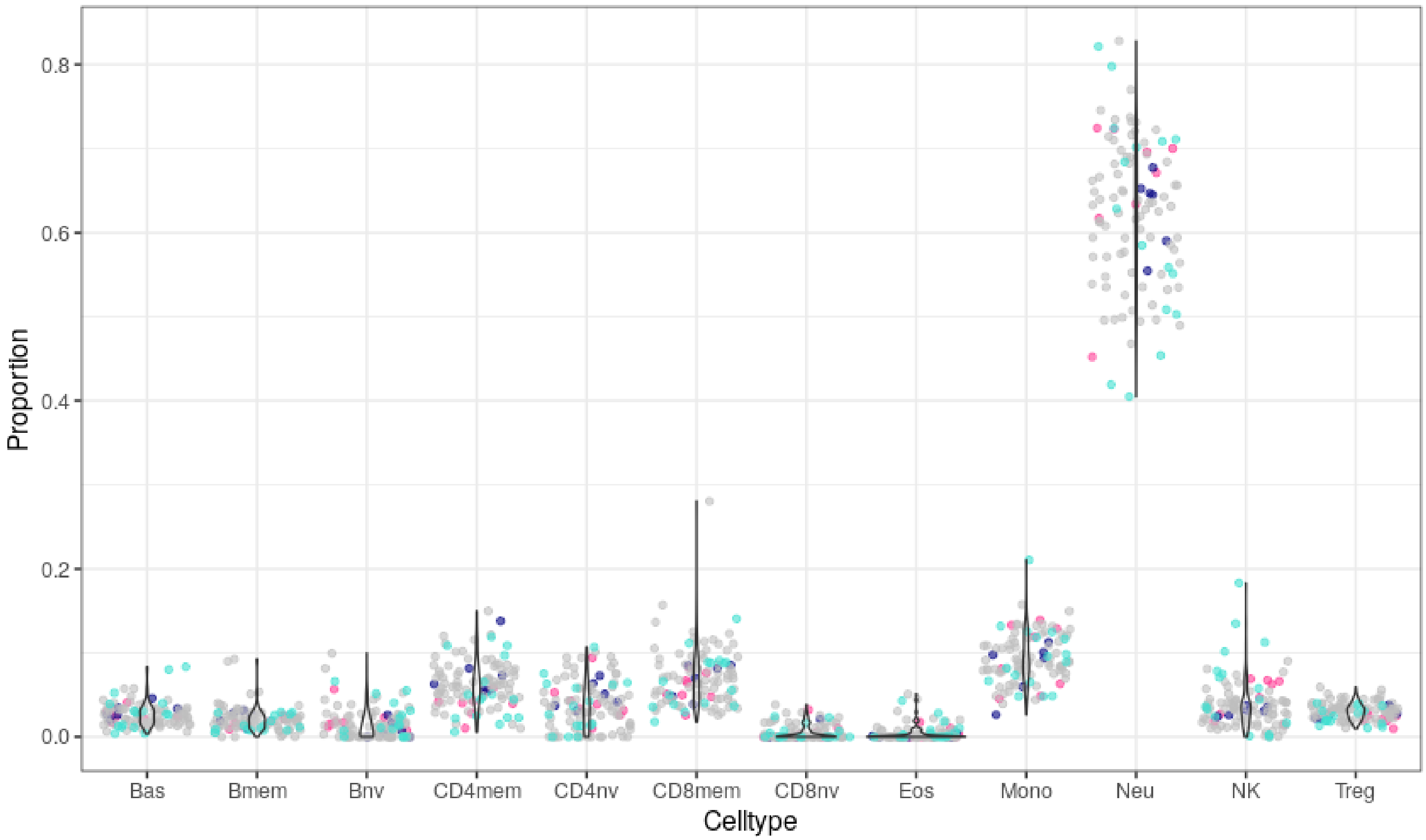


***Figure S3 Estimated blood cell type proportions.*** *Twelve blood cell type proportions in peripheral blood were estimated based on epigenomic deconvolution. Different colours indicate the different sub-datasets present in the superset.*

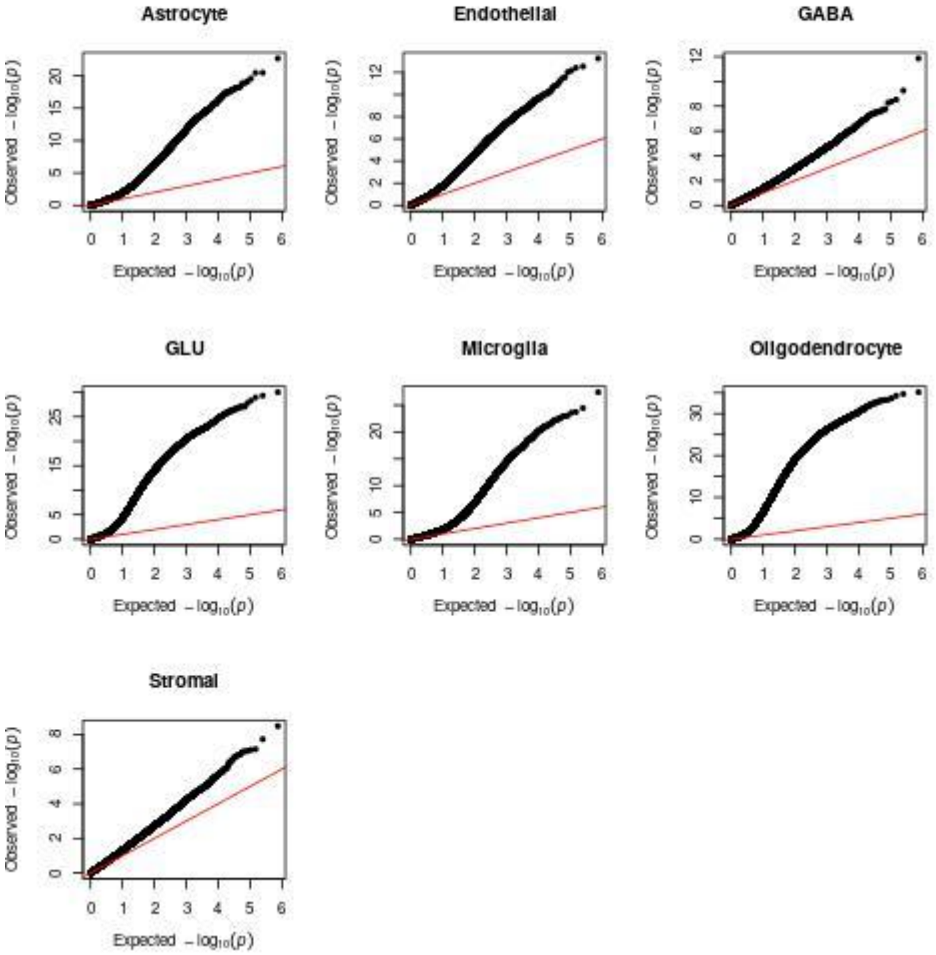


***Figure S4 Quantile-quantile plots for meta-analysis of epigenome-wide association studies in brain tissue of cell type proportions in the frontal brain.*** *Expected -$\log_{10}$(p-values) on the x-axis are plotted against the observed -$\log_{10}$(p-values) on the y-axis. Lambda values for the different meta-epigenome-wide association studies are the following: Astrocyte = 1.89; Endothelial = 1.80; GABA = 1.46; GLU = 3.03; Microglia = 1.76; Oligodendrocyte = 4.23; Stromal = 1.42*

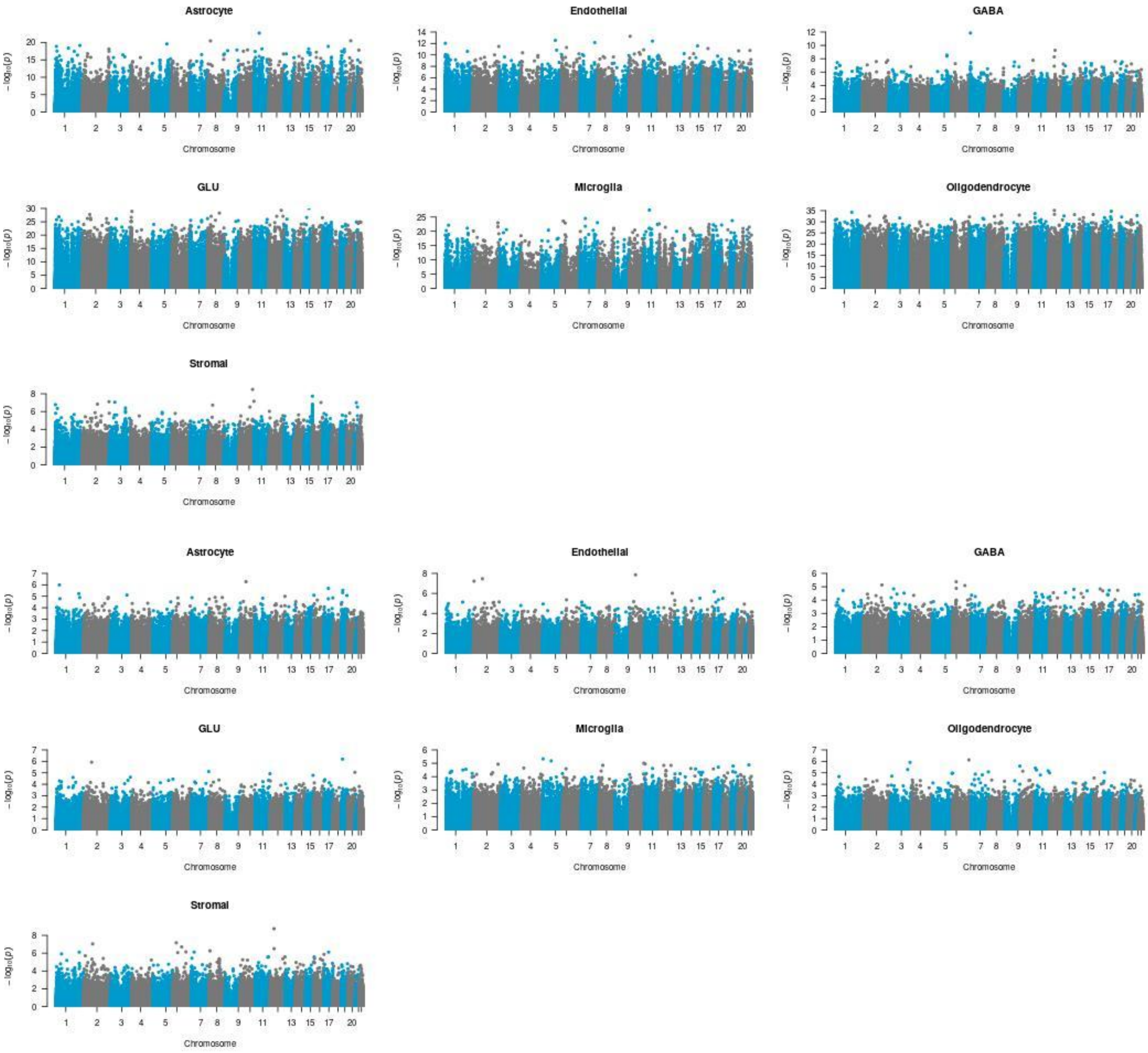


***Figure S5 Manhattan plots of meta-analysis of epigenome-wide association studies for brain cell types in the frontal cortex and blood.*** *Every panel depicts the meta-analysis result of one brain cell type. The -log$_{10}$(p-values) of the DNA methylation sites (y-axis) are plotted along their chromosomal position(x-axis). The upper panels reflect analyses performed in the frontal cortex, whereas the lower panels reflect analyses performed in peripheral blood.*

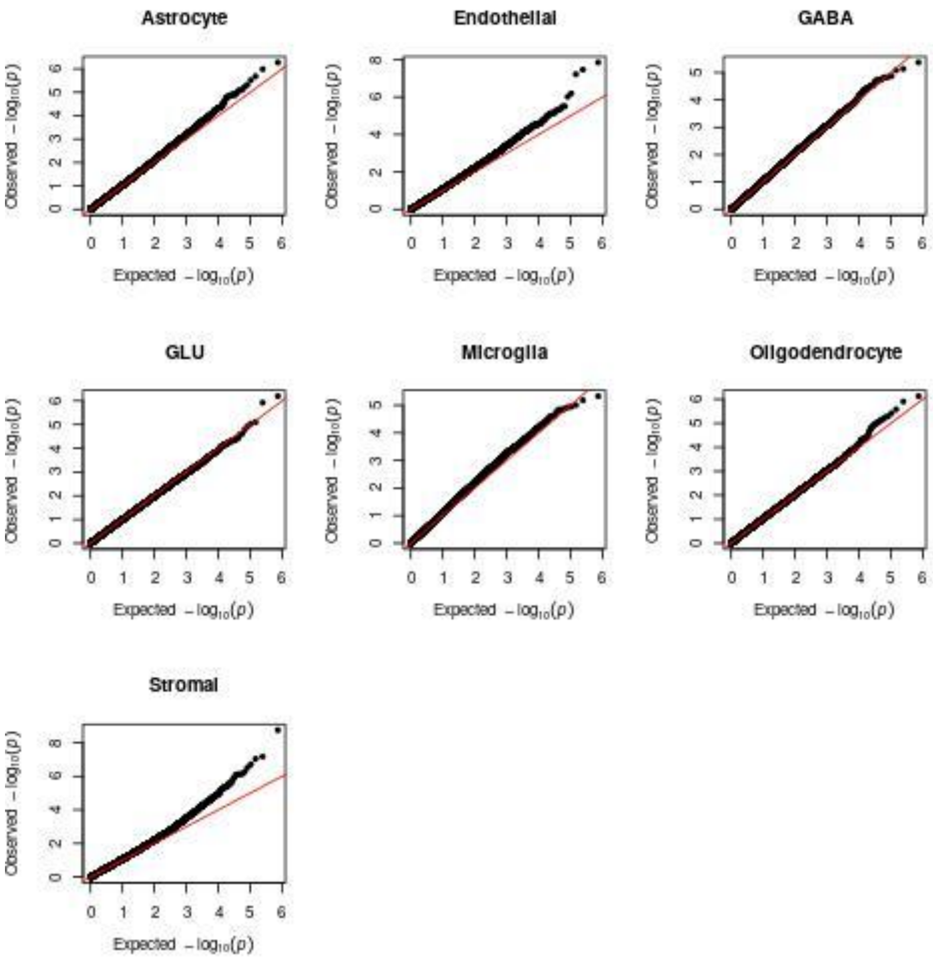


***Figure S6 Quantile-quantile plots for meta-analysis of epigenome-wide association studies in blood of cell type proportions in the frontal brain.*** *Expected -log$_{10}$(p-values) on the x-axis are plotted against the observed -log$_{10}$(p-values) on the y-axis. Lambda values for the different meta-epigenome-wide association studies are the following: Astrocyte = 1.03; Endothelial = 1.09; GABA = 1.04; GLU = 1.00; Microglia = 1.15; Oligodendrocyte = 1.05; Stromal = 1.06*

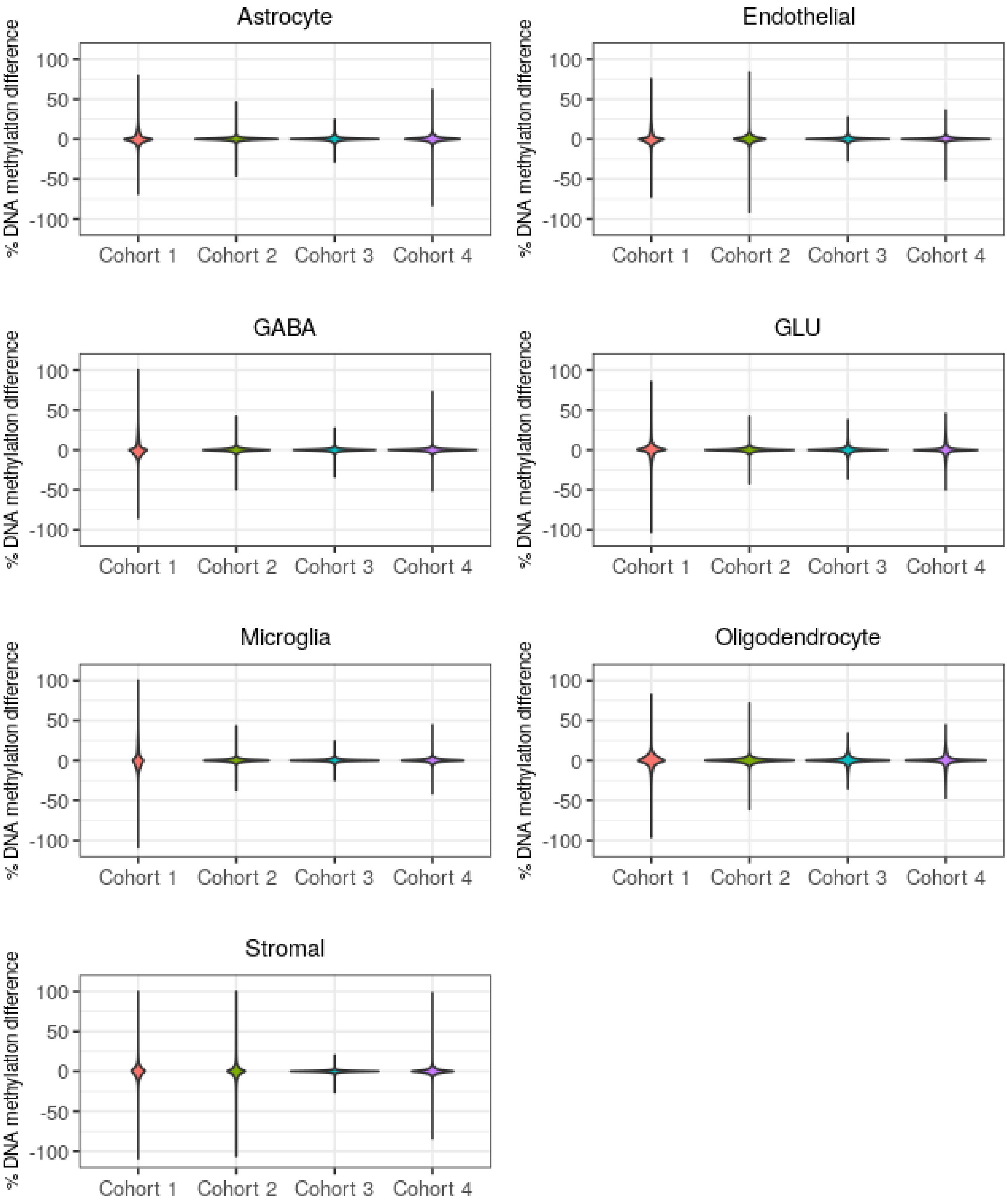


***Figure S7 Effect sizes of epigenome-wide association studies are similar for each cell type proportion in the frontal brain.*** *The percentage DNA methylation differences between the sample with the highest and lowest cell type proportions was calculated by multiplying the EWAS regression coefficient of the cell type of interest with the range of that cell type proportions between the 5th and 95th percentile to reduce the effects of outliers. The delta betas are shown for the EWAS in each cohort separately.*

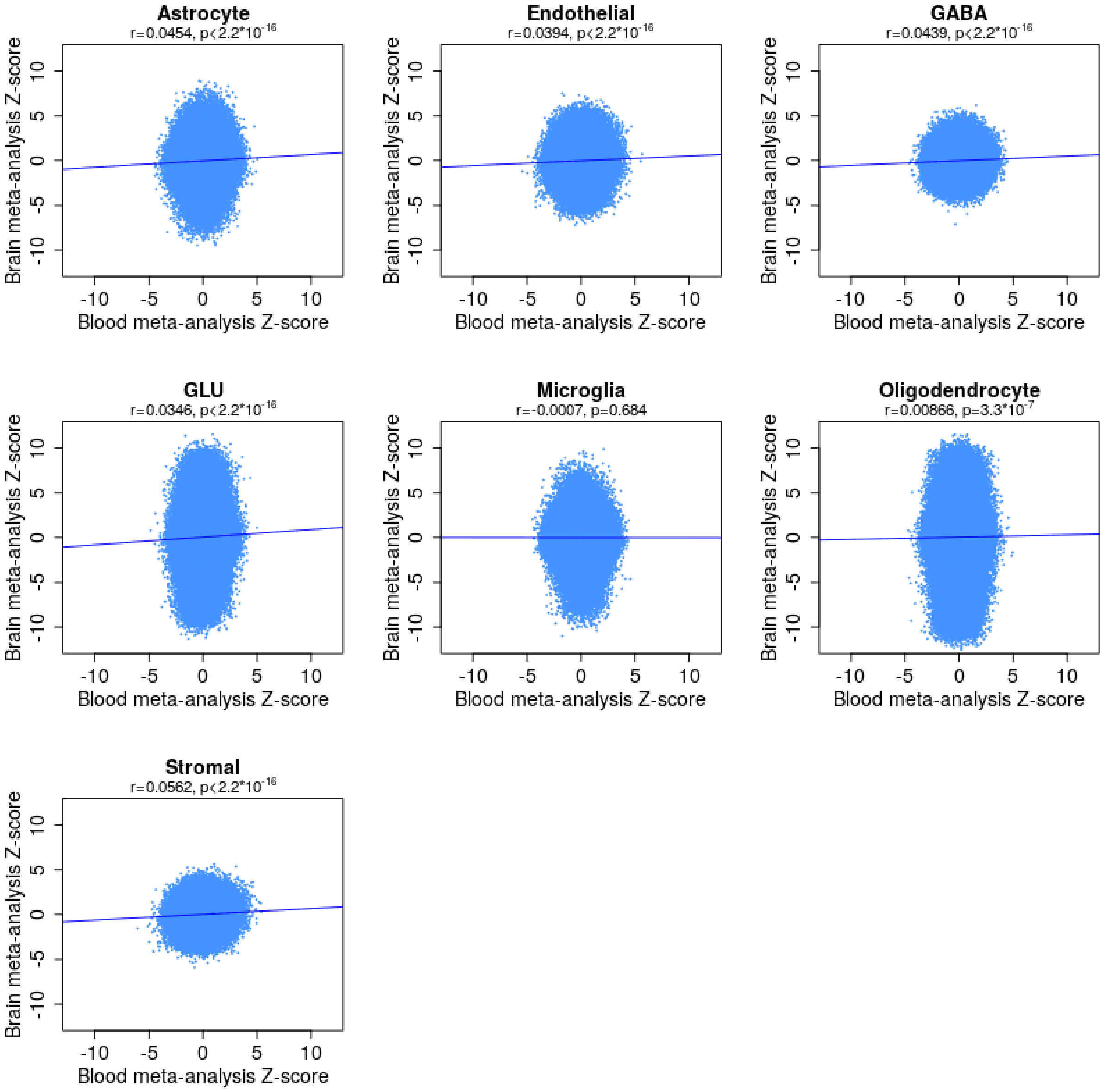


*Figure S8 Concordance between Z-scores from meta-analysis of epigenome-wide association studies for brain cell types in the frontal brain and blood. Effect sizes (Z-scores) of the meta-analyses for brain cell types in blood (x-axis) are plotted against the effect sizes (Z-scores) of the meta-analyses for brain cell types in the frontal cortex. Each dot represents on CpG site. Pearson's correlation coefficients (r) and corresponding p-values (p) are reported for the Z-scores for each brain cell type.*

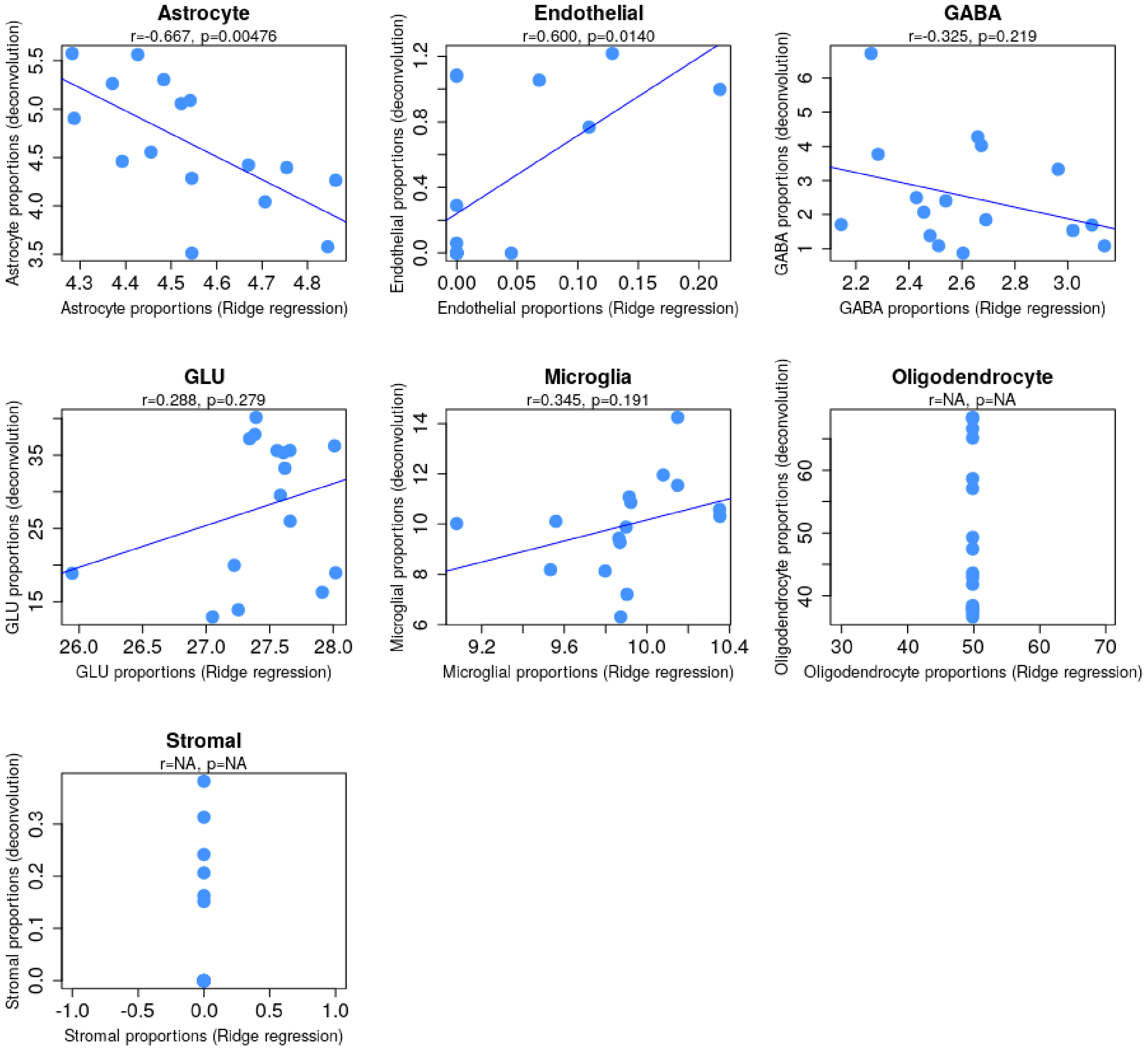


***Figure S9 Ridge regression performance for brain cell type proportion estimation based on Variable Methylation Regions in whole blood DNA methylation.*** *Estimated brain cell type proportions based on epigenomic deconvolution in DNA methylation of frontal cortex samples is plotted on the y-axis. Brain cell type proportions estimated based on the optimal models from Ridge regression are plotted the x-axis. Every dot represents a single sample from the renormalized train datasets 4 (**Table S1**). The Pearson's correlation (r) and p-value (p) are given for every brain cell type.*

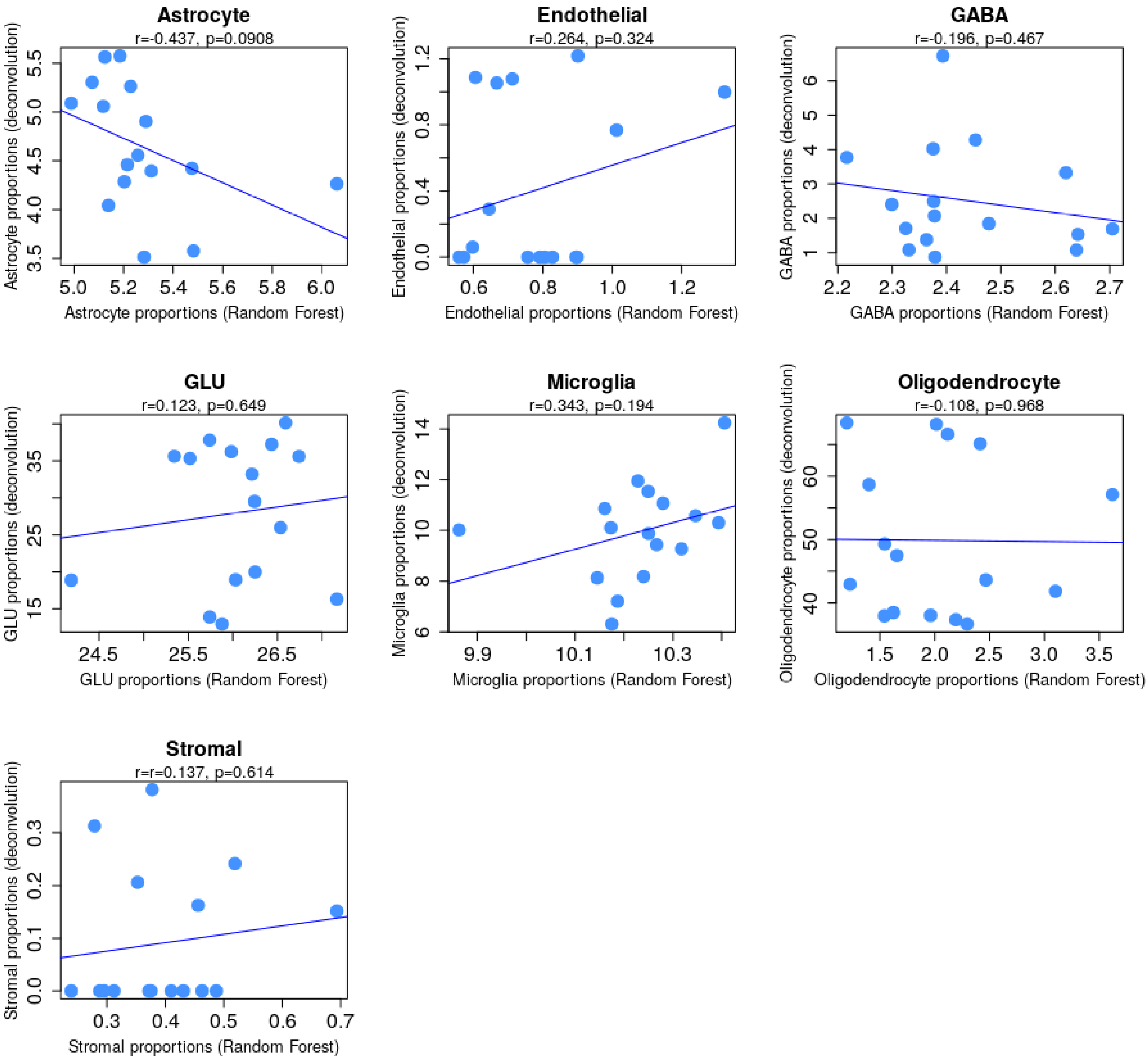


***Figure S10 Random Forest performance for brain cell type proportion estimation based on Variable Methylation Regions in whole blood DNA methylation.*** *Estimated brain cell type proportions based on epigenomic deconvolution in DNA methylation of frontal cortex samples is plotted on the y-axis. Brain cell type proportions estimated based on the optimal models from Random Forest are plotted the x-axis. Every dot represents a single sample from the renormalized train datasets 4 (**Table S1**). The Pearson's correlation (r) and p-value (p) are given for every brain cell type.*

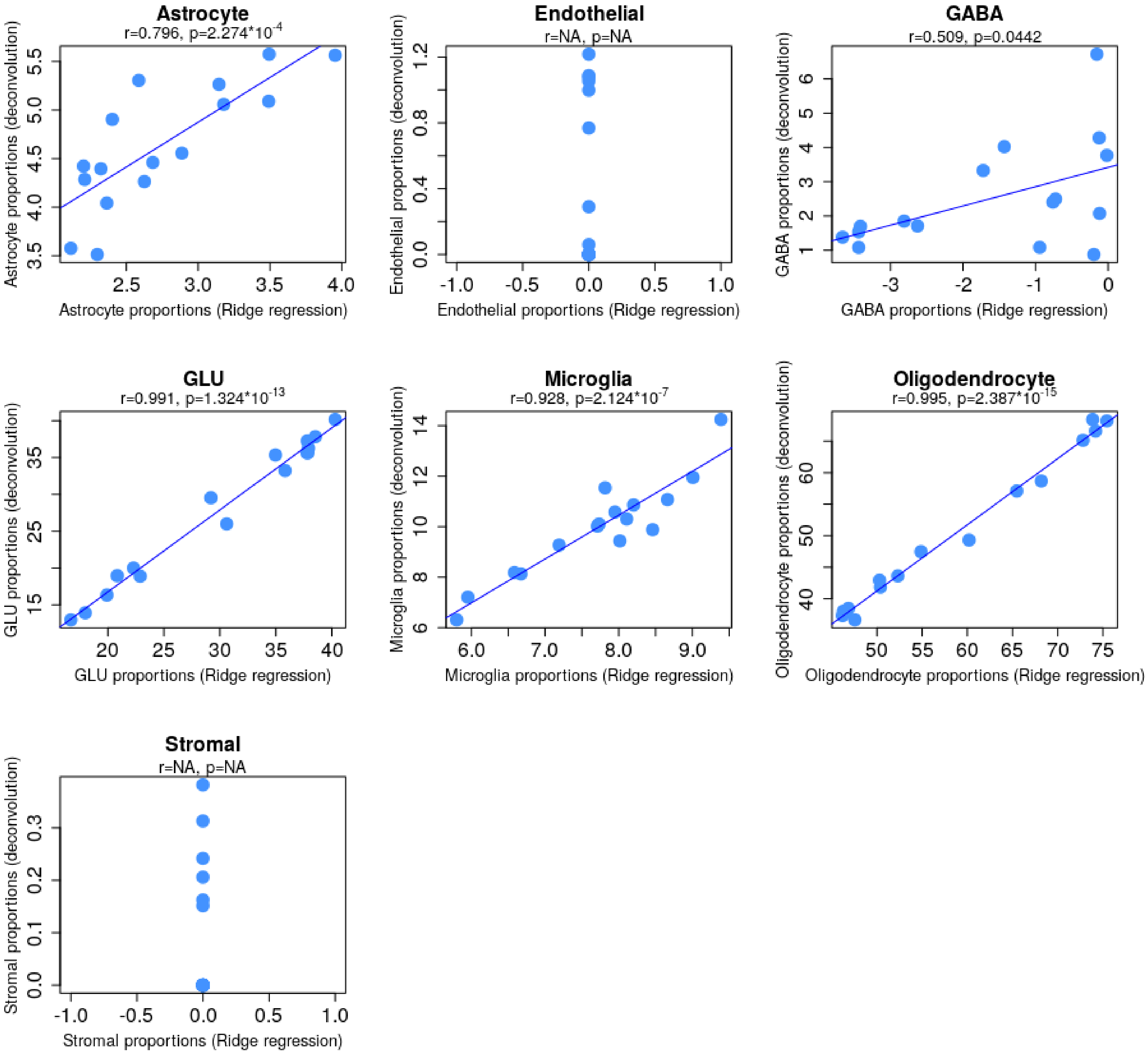


*Figure S11 Ridge regression performance for brain cell type proportion estimation based on frontal cortex DNA* ***methylation.*** *Estimated brain cell type proportions based on epigenomic deconvolution in DNA methylation of frontal cortex samples is plotted on the y-axis. Brain cell type proportions estimated based on the optimal models from Ridge regression are plotted the x-axis. Every dot represents a single sample from the renormalized train datasets 4 (**Table S1**). The Pearson's correlation (r) and p-value (p) are given for every brain cell type.*

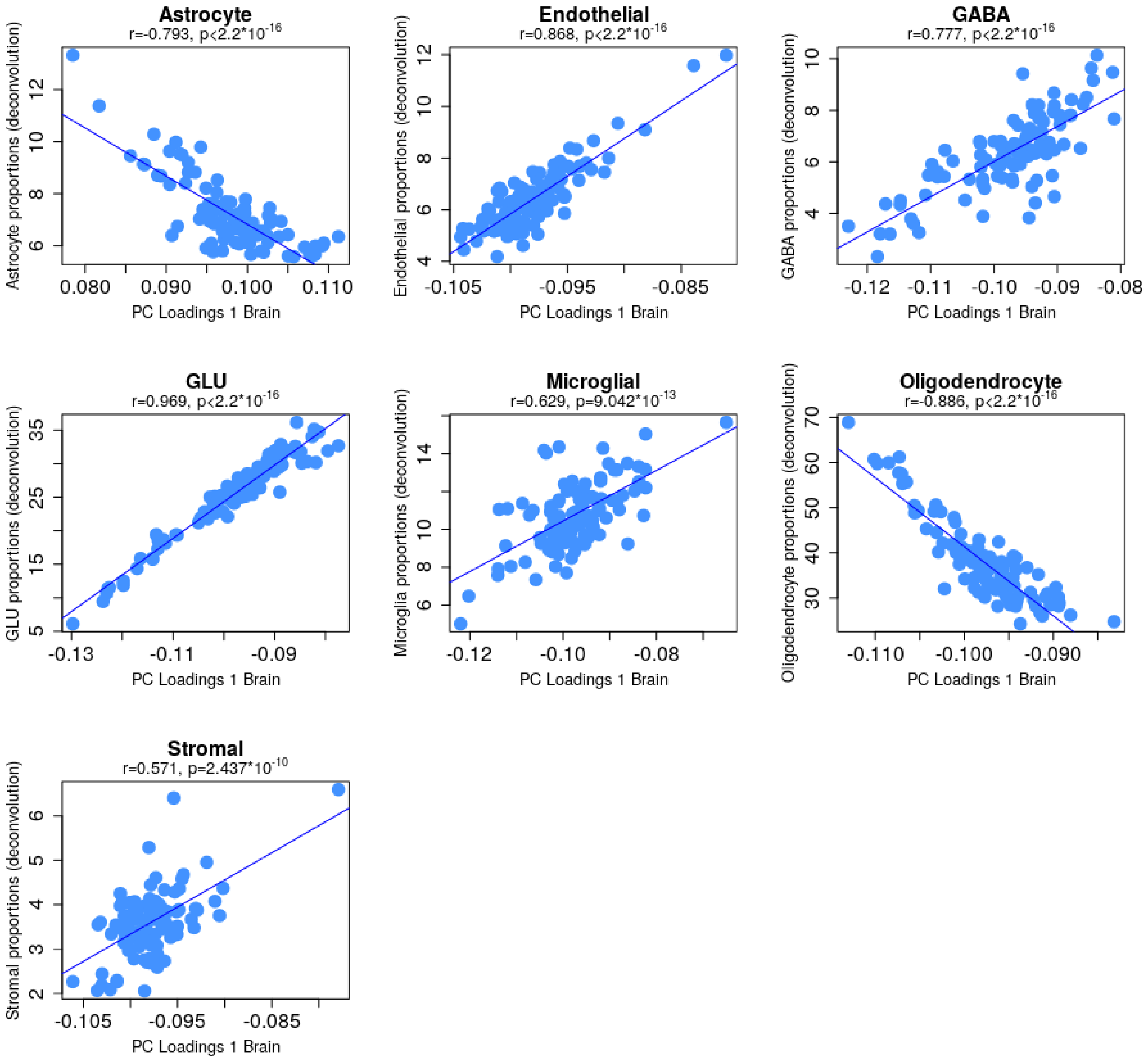


***Figure S12 DNAm sites in the frontal cortex with high blood-brain correlation are associated with brain cell type proportion in the frontal brain.*** *Estimated brain cell type proportions based on epigenomic deconvolution in DNA methylation of frontal cortex samples is plotted on the y-axis. The loadings of the first principal component (PC) on DNAm in the brain based on the highly blood-brain correlating probes in the reference panel for the specific brain cell types are plotted on the x-axis. Every dot represents a single sample form the combined matched blood-brain dataset. The Pearson's correlation (r) and p-value (p) are given for every brain cell type.*

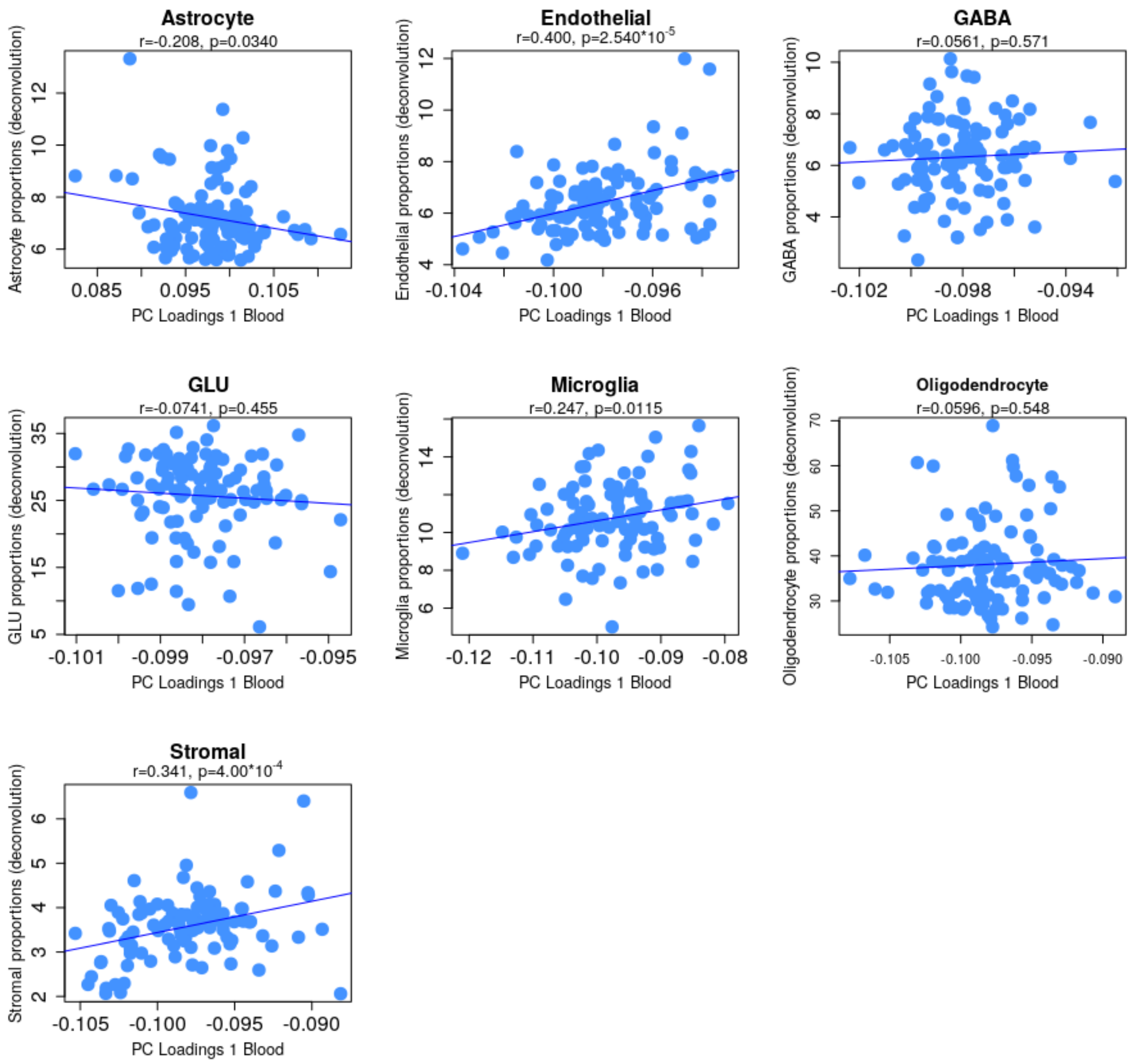


***Figure S13 DNAm sites in whole blood with high blood-brain correlation are associated with brain cell type proportion in the frontal brain.*** *Estimated brain cell type proportions based on epigenomic deconvolution in DNA methylation of frontal cortex samples is plotted on the y-axis. The loadings of the first principal component (PC) on DNAm in whole blood based on the highly blood-brain correlating probes in the reference panel for the specific brain cell types are plotted on the x-axis. Every dot represents a single sample form the combined matched blood-brain dataset. The Pearson's correlation (r) and p-value (p) are given for every brain cell type.*

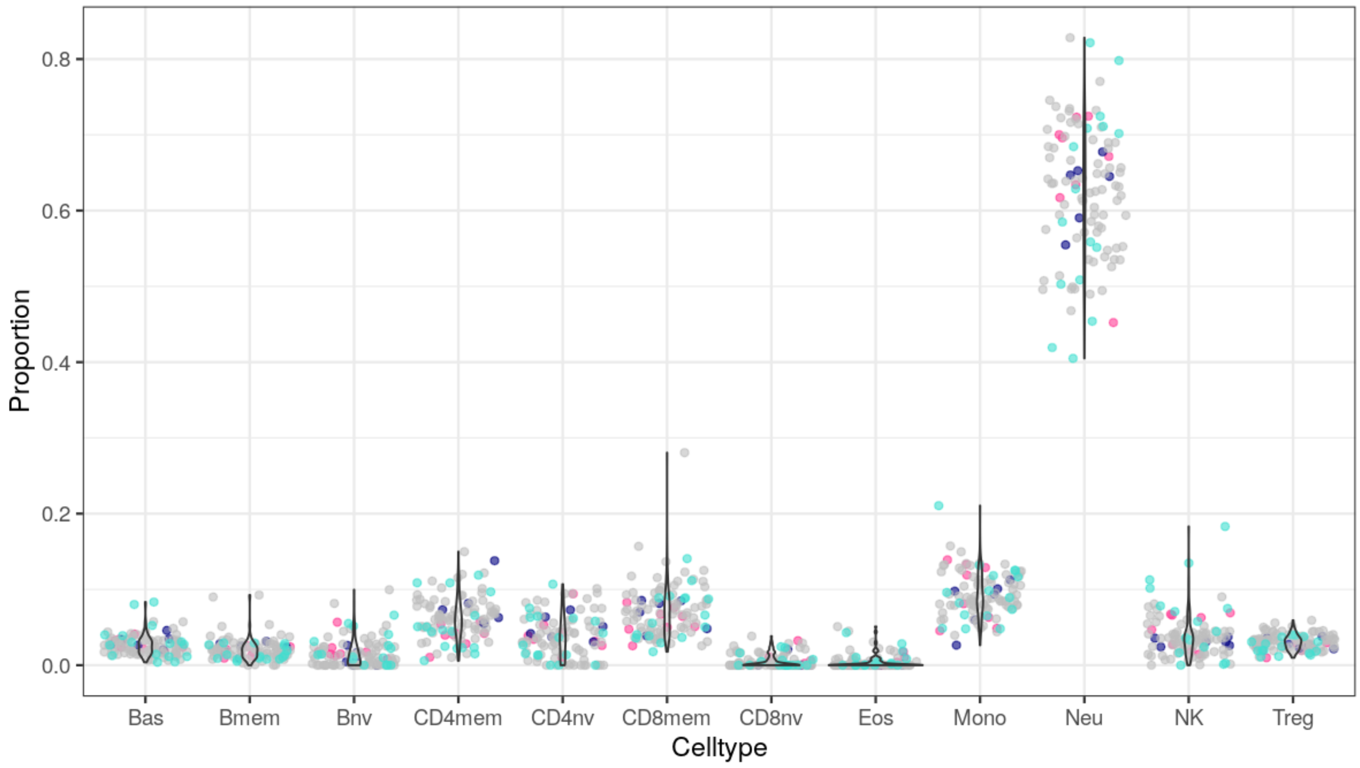


***Figure S14 Estimated blood cell type proportions.*** *Blood cell type proportions in whole blood estimated based on epigenomic deconvolution with an enhanced reference panel for twelve different cell types. Different colours indicate the different sub-datasets present in the combined dataset. Bas=Basophils; Bmem = B memory cells; Bnv = naive B cells; CD4mem = CD4*$^{+}$ *T memory cells; CD4nv = naive CD4*$^{+}$ *T cells; CD8mem = CD8*$^{+}$ *T memory cells; CD8nv = naive CD8*$^{+}$ *T cells; Eos = Eosinophils; Mono = Monocytes; Neu = Neutrophils; NK= Natural Killer cells; Treg = Regulatory T cells.*

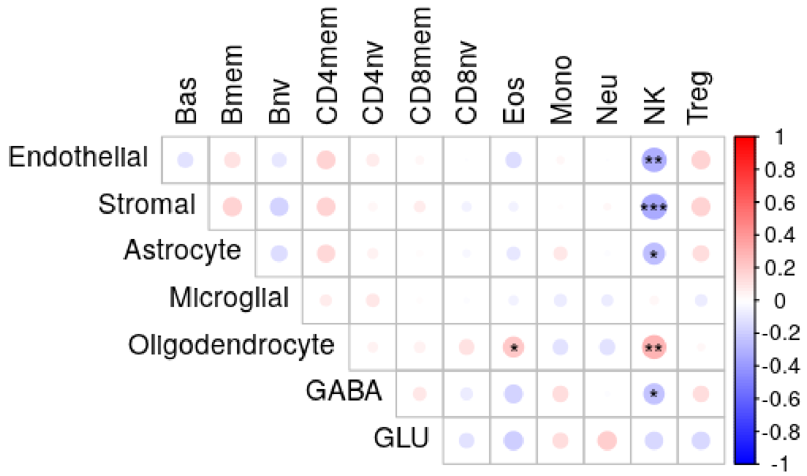


***Figure S15 Correlation between estimated blood and brain cell types.*** *Pearson's correlations were calculated between brain cell types (y-axis) and blood cell types (x-axis) from matched individuals, based on epigenomic deconvolution estimates. Size and colour of the circles indicate size of the Pearson's correlation (bright red: Pearson's r = 1, white: Pearson's r=0, dark blue: Pearson's r=-1). *p-value<0.05, **p-value<0.01, ***p-value<0.001*

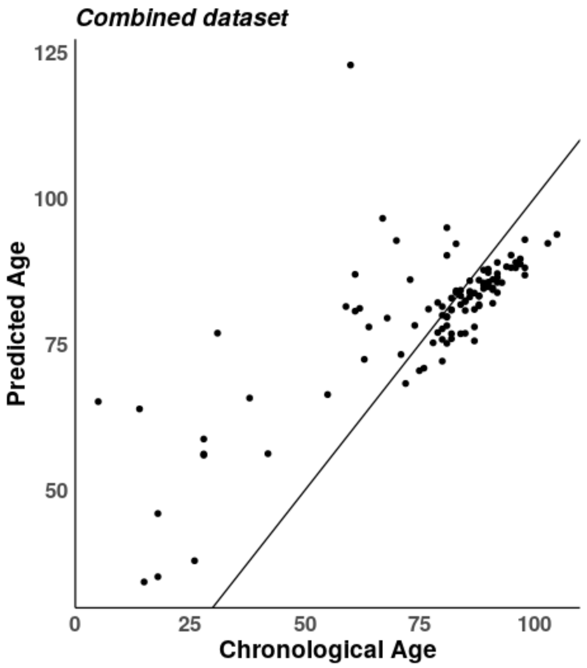


***Figure S16 Associations between chronological and biological age.*** *Chronological age is plotted against the x-axis. On the y-axis, the predicted age based on the cortical clock is plotted.*

### References

1. Aryee MJ, Jaffe AE, Corrada-Bravo H, Ladd-Acosta C, Feinberg AP, Hansen KD *et al.* Minfi: a flexible and comprehensive Bioconductor package for the analysis of Infinium DNA methylation microarrays. *Bioinformatics* 2014; **30**(10)**:** 1363-1369.

2. Pidsley R, Zotenko E, Peters TJ, Lawrence MG, Risbridger GP, Molloy P *et al.* Critical evaluation of the Illumina MethylationEPIC BeadChip microarray for whole-genome DNA methylation profiling. *Genome Biol* 2016; **17**(1)**:** 208.

3. McCartney DL, Walker RM, Morris SW, McIntosh AM, Porteous DJ, Evans KL. Identification of polymorphic and off-target probe binding sites on the Illumina Infinium MethylationEPIC BeadChip. *Genom Data* 2016; **9:** 22-24.

4. Pidsley R, CC YW, Volta M, Lunnon K, Mill J, Schalkwyk LC. A data-driven approach to preprocessing Illumina 450K methylation array data. *BMC Genomics* 2013; **14:** 293.

5. Johnson WE, Li C, Rabinovic A. Adjusting batch effects in microarray expression data using empirical Bayes methods. *Biostatistics* 2007; **8**(1)**:** 118-127.

6. Friedman JH, Hastie, T., & Tibshirani, R. Regularization Paths for Generalized Linear Models via Coordinate Descent. *Journal of Statistical Software* 2010; **33**(1)**:** 1-22.

7. Tay JK, Narasimhan, B., & Hastie, T. . Elastic Net Regularization Paths for All Generalized Linear Models. *Journal of Statistical Software* 2023; **106**(1)**:** 1-31.

8. Gatev E, Gladish N, Mostafavi S, Kobor MS. CoMeBack: DNA methylation array data analysis for co-methylated regions. *Bioinformatics* 2020; **36**(9)**:** 2675-2683.

9. Shireby GL, Davies JP, Francis PT, Burrage J, Walker EM, Neilson GWA *et al.* Recalibrating the epigenetic clock: implications for assessing biological age in the human cortex. *Brain* 2020; **143**(12)**:** 3763-3775.

10. Zhang Q, Vallerga CL, Walker RM, Lin T, Henders AK, Montgomery GW *et al.* Improved precision of epigenetic clock estimates across tissues and its implication for biological ageing. *Genome Med* 2019; **11**(1)**:** 54.

11. Horvath S. DNA methylation age of human tissues and cell types. *Genome Biol* 2013; **14**(10)**:** R115.
